# High air humidity dampens salicylic acid pathway and plant resistance via targeting of NPR1

**DOI:** 10.1101/2022.10.28.514180

**Authors:** Lingya Yao, Zeyu Jiang, Yiping Wang, Shiwei Wan, Xiu-Fang Xin

## Abstract

The occurrence of plant disease is determined by interactions among host, pathogen and climate conditions. Air humidity has long been recognized to profoundly influence diseases in the phyllosphere and high air humidity (e.g., after rain falls) is known as a prerequisite for numerous disease outbreaks in the field^1–3^. However, the molecular basis of how high humidity interferes with plant resistance mechanisms to favor disease remained elusive. Here we show that high humidity is associated with an “immune-compromised” status of plants, revealed by lower expression of defense genes during bacterial infection of Arabidopsis plants. Examination of humidity’s effect on individual immune pathways showed that the accumulation and signaling of salicylic acid (SA), an essential hormone conferring plant resistance against infectious microbes^4,5^, are significantly inhibited under high humidity. Surprisingly, NPR1 protein, an SA receptor and central transcriptional co-activator of SA-responsive genes^6–9^, accumulated to a significantly higher level in the nucleus under high humidity. Further investigation indicated a decreased binding affinity of NPR1 protein to the target gene promoter, suggestive of an “inactive” nature of NPR1, under high humidity and an impaired ubiquitination and degradation of NPR1 protein, likely due to down-regulation of Cullin 3-mediated cellular ubiquitination pathway and 26S proteasome pathway under high humidity. Our study uncovers disruption of NPR1 protein turnover as a major mechanism, by which high humidity dampens plant immune strength against pathogens, and provides new insights into the long-observed air humidity influence on diseases in nature.

## Main

The “plant-pathogen-environment” interaction is a fundamental principle for understanding disease epidemics^10,11^. Over the past decades, a great amount of efforts have been devoted to unravel the convoluted interplay between plant immunity and pathogen virulence. In contrast, the mechanistic understanding of the environmental influence on disease development is rather limited. Plants perceive danger signals from pathogens, through cell surface receptors and intracellular receptors, to activate downstream signaling cascades called pattern triggered immunity (PTI) and effector triggered immunity (ETI)^12–14^. Pathogen recognition further induces defense-related hormone pathways, such as SA, jasmonic acid (JA) and ethylene pathways, which amplify immune responses and enhance resistance against pathogens^15^.

It is well documented that high air humidity, a condition typically occurring after rain falls or in tropical or coastal regions, strongly promotes a variety of diseases caused by bacteria, fungi or oomycete in the aerial parts of the plant in agricultural ecosystems^1–3,16^. Previous studies showed that high humidity promotes the virulence of bacterial pathogens such as *Pseudomonas syringae*, by facilitating the generation of a water-rich living environment in the leaf tissue^17–19^, and is important for fungal spores to germinate^20^. High humidity also impedes stomatal closure and ETI-associated cell death in plants^21,22^, and daily humidity oscillation improves plant fitness-related traits and enhances ETI at night^23^. Despite of these progress, how high humidity affects plant immunity was not known at the mechanistic level.

## Impairment of plant immunity under high humidity

To investigate humidity effect on plant responses, we set up an experimental system in which plants were first grown under ambient humidity (i.e., 60% relative humidity, RH) till four weeks old and then placed under high humidity (~95% RH), to simulate the humidity condition that usually forecasts disease outbreaks in nature^3,16^, or low humidity (~45% RH). Plants appeared to grow bigger (based on rosette size) with a higher fresh weight, but no change in dry weight, several days after high humidity treatment (Extended Data Fig. 1). Under this setting, high humidity strongly promoted diseases caused by *P. syringae* pv. *tomato* (*Pst*) DC3000 bacteria in Arabidopsis leaves (Fig. 1a-b). We then used this system to study whether high humidity affects plant defense. The Col-0 plants were treated with different humidity and infiltrated with *Pst* DC3000 bacteria, and the expression of immune-related genes was examined. We found that the transcriptional induction of defense responsive genes, such as *WRKY29, NHL10, FRK1, PR2, EDS5* and *EDS1*, was dramatically suppressed under high humidity (Fig. 1c), suggesting that high humidity indeed influences plant defense and places plants to an “immune-compromised” condition, presumably contributing to severity of diseases.

**Fig. 1.**
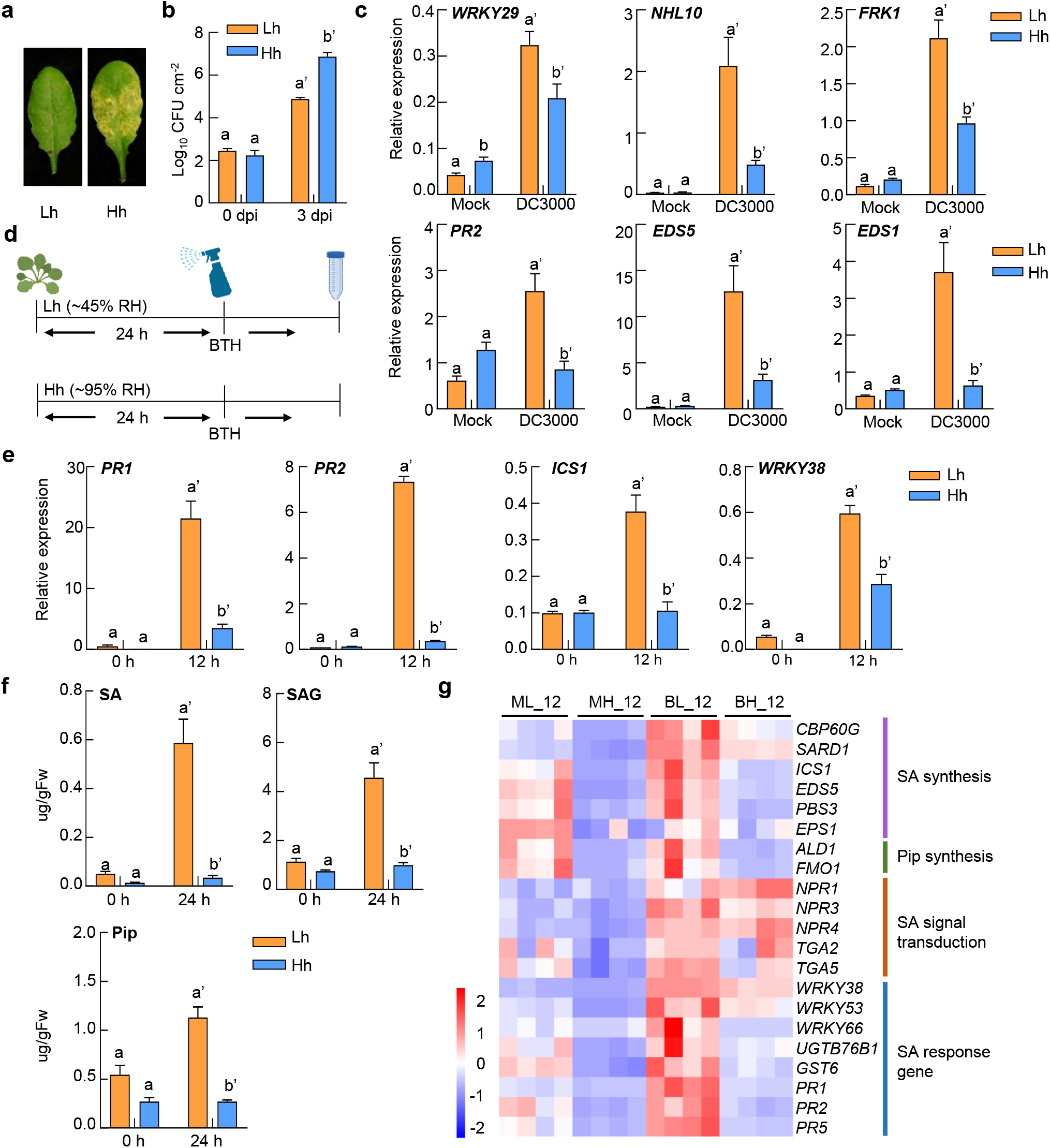
Plant defense-related gene expression and SA pathway are suppressed under high humidity in Arabidopsis. **a-b**, High humidity strongly promotes *P. syringae* infection in Arabidopsis. Four-week-old Col-0 plants (grown under ~60% RH) were infiltrated with *Pst* DC3000 at 1×10^6^ cfu/ml and placed under low humidity (Lh, ~45% RH) or high humidity (Hh, ~95% RH). Disease symptoms (**a**) and bacterial population (**b**) were recorded 3 days later. Data represent mean ± standard error of the mean (SEM) (n = 3 biological replicates). Different letters indicate statistically significant differences, as analyzed by two-way ANOVA with Tukey’s test (p<0.05). **c**, RT-qPCR analysis of *WRKY29, NHL10, FRK1, PR2, EDS5* and *EDS1* transcript levels in Col-0 plants under low or high humidity. Col-0 plants (grown under ~60% RH) were infiltrated with *Pst* DC3000 at 1×10^8^ cfu/ml and placed under different humidity. Leaf tissues were sampled 5 h after infiltration for RNA extraction and RT-qPCR. Data represent mean ± SEM (n = 4 biological replicates). Different letters indicate statistically significant differences, as analyzed by two-way ANOVA with Tukey’s test (p<0.05). **d**, A schematic diagram showing the humidity and BTH treatment on plants for results in **e-g**. Four-week-old Col-0 plants grown under ambient humidity (60% RH) were pre-treated with high or low humidity for 24 h, sprayed with100 μM BTH and placed back to different humidity setting before sampling. **e**, RT-qPCR analysis of *PR1*, *PR2*, *ICS1* and *WRKY38* expression levels in plants under Lh or Hh, at 0 h and 12 h after BTH treatment. Data represent mean ± SEM (n = 3 biological replicates). Different letters indicate statistically significant differences, as analyzed by two-way ANOVA with Tukey’s test (p<0.05). **f**, Quantification of SA, SAG and Pip levels under Lh or Hh, at 0 h and 24 h after BTH treatment. Data represent mean ± SEM (n = 4 biological replicates). Different letters indicate statistically significant differences, as analyzed by two-way ANOVA with Tukey’s test (p<0.05). Experiments were repeated three times with similar results. **g**, A heatmap of SA pathway-related genes under Lh or Hh, at 12 h after BTH/mock treatment, in the RNAseq results. ML_12, 12 h after mock treatment under low humidity; MH_12, 12 h after mock treatment under high humidity; BL_12, 12 h after BTH treatment under low humidity; BH_12, 12 h after BTH treatment under high humidity.

## Inconsistent effects of high humidity on PTI or JA pathway

Infection of pathogenic microorganisms like *Pst* DC3000 simultaneously activates multiple plant immune signaling pathways, including PTI, SA, JA and ethylene pathways. Furthermore, these pathways show extensive crosstalk with each other^15,24^, making it difficult to clearly dissect humidity influence on specific pathways. We therefore applied elicitors to induce individual pathways and analyzed which immune pathway(s) is altered under high humidity. As shown in Extended Data Fig. 2a, plants were treated with flg22, a bacterial flagellin-derived peptide inducing PTI^25^, and PTI responses under different humidity settings were examined. We observed an increase in the expression of flg22-responsive genes, such as *WRKY29* and *IOS1*, but similar MPK3/6 phosphorylation and ROS production under high humidity (Extended Data Fig. 2b-d). We next examined whether humidity affects the JA pathway, another important defense hormone pathway^26^. We found that, while the MeJA-induced accumulation of JA and JA-Ile, the bioactive form of the hormone, was reduced under high humidity, the expression of MeJA-induced genes such as *JAZ8, PDF1.2, LOX2* and *VSP2* show different or opposite regulation patterns under high humidity (Extended Data Fig. 3). Overall, these data indicate that high humidity affects different PTI and JA responses in different manners.

## Significant inhibition of SA pathway under high humidity

SA is a beta-hydroxy phenolic acid and plays an essential role in resistance against a variety of bacterial, fungal or viral pathogens in plants^5,27,28^. SA pathway can be activated by the treatment of benzothiadiazole (BTH), a SA analog without toxicity to plant cells. We found that the expression of BTH-induced SA response genes, including *PR1*, *PR2*, *ICS1* and *WRKY38*, was consistently and significantly lower under high humidity (Fig. 1d-e). In addition, BTH-induced accumulation of SA, salicylic acid beta-glucoside (SAG) and pipecolic acid (Pip), which is synergistic to SA in activating plant defense^29^, was strongly suppressed under high humidity (Fig. 1f).

To investigate the effect of high humidity on the Arabidopsis transcriptome and whether high humidity targets specific branches of the SA pathway, we performed an RNAseq experiment. Col-0 plants were pre-treated with high or low humidity for 24 h, sprayed with BTH (water as control) and sampled 12 h later for total RNA extraction and sequencing (Fig. 1d). We found that, high humidity led to 4005 and 2515 differentially-regulated genes (fold change>2; adjusted p value<0.01), under mock and BTH treatment, respectively (Extended Data Fig. 4c-d). Gene Ontology analysis showed that “response to bacteria, SA, water deprivation or temperature”-related pathways were enriched in humidity-regulated genes (Extended Data Fig. 4e-f). Notably, about ~50% (474/927) of BTH-responsive genes showed differential expression under different humidity settings (Extended Data Fig. 4b), suggesting that high humidity broadly affects SA responses. Furthermore, both basal and BTH-induced expression of many SA/Pip biosynthesis-related genes, including *ICS1, CBP60g, PBS3, ALD1* and *FMO1*, and SA-responsive marker genes, including *PR1*, *PR2* and *PR5*, was significantly inhibited under high humidity (Fig. 1g). Genes involved in SA signal transduction, such as *NPR1, NPR3/4* and *TGA2*, were not transcriptionally regulated by humidity under our experimental conditions.

To determine the biological relevance of high humidity suppression of SA pathway, we performed *Pst* bacterial infection assay under different humidity in the *sid2-2* mutant^30^, which lacks the key SA biosynthesis enzyme -isochorismate synthase 1 (ICS1), and *npr1-6* mutant^31^, which lacks the SA receptor NPR1. While *Pst* DC3000 grew poorly and did not cause visible disease symptoms on Col-0 plants under low humidity, it caused disease symptoms and multiplied significantly higher in the *sid2-2* and *npr1-6* plants under low humidity, almost to the level observed in Col-0 plant under high humidity (Fig. 2a-b). This disease restoration under low humidity was not obvious in the PTI receptor/co-receptor mutant plant, *fls2/efr/cerk1*^17^, which echoes the insignificant effect of high humidity on PTI pathway. However, both *sid2-2* and *npr1-6* mutations also increased plant susceptibility to *Pst* infection under high humidity, albeit to a lesser degree than under low humidity, likely due to high humidity effect on other disease-related processes such as pathogen virulence promotion^17^. Altogether, these results suggest that suppression of SA pathway is a significant determinant of enhanced Arabidopsis susceptibility to *Pst* infection under high humidity.

**Fig. 2.**
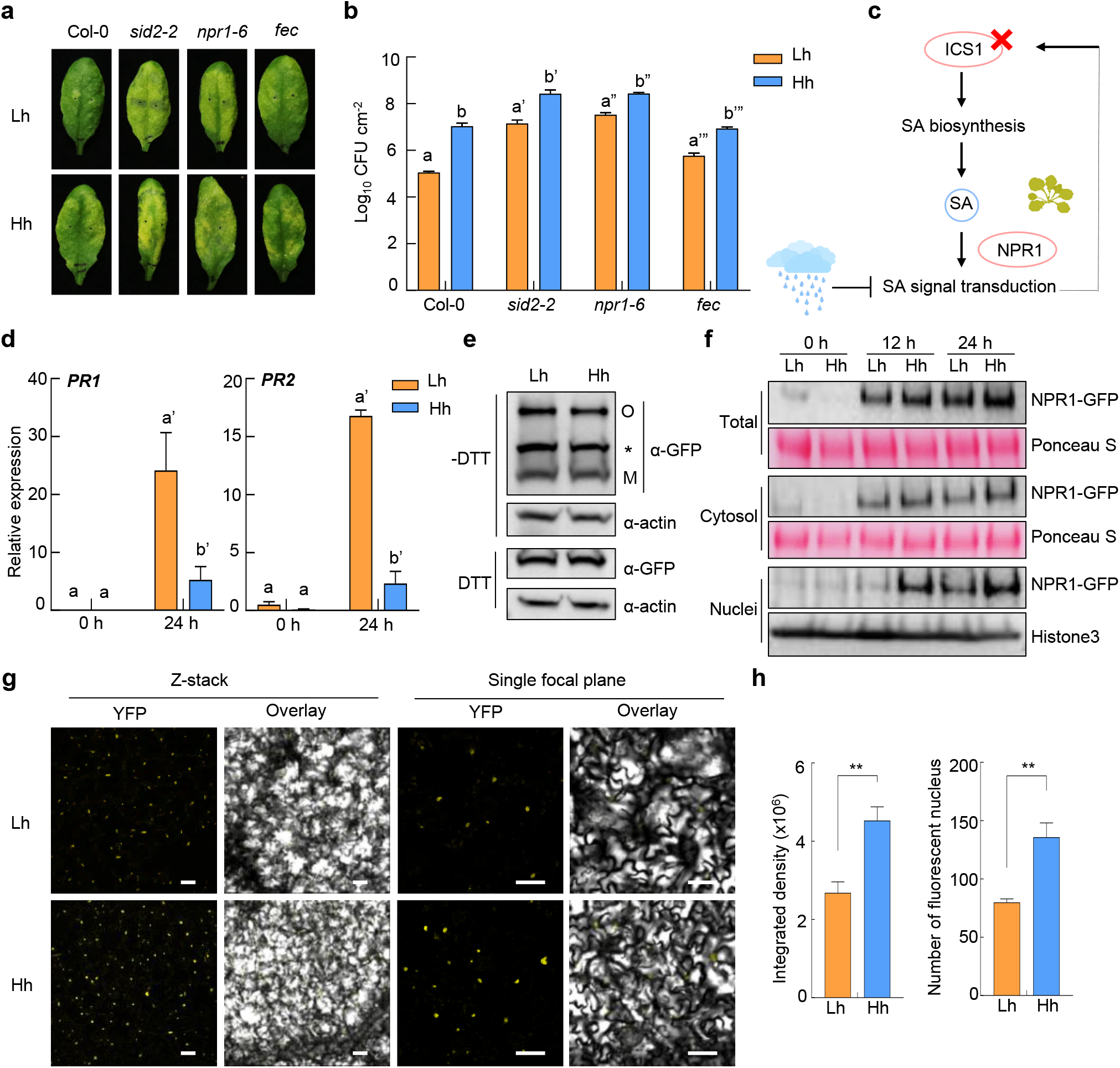
NPR1 protein is over-accumulated in the nucleus of the plant cell under high humidity. **a-b**, SA pathway mutation partially rescued disease susceptibility under low humidity. Col-0, *sid2-2, npr1-6* and *fec* plants were pre-treatment with low or high humidity for 24 h, infiltrated with *Pst* DC3000 at 1×10^6^ cfu/ml and placed back to different humidity. Disease symptoms (**a**) and bacterial population (**b**) were recorded 3 days post infiltration. Data in **b** represent mean ± SEM (n = 3 biological replicates). Different letters indicate statistically significant differences, as analyzed by two-way ANOVA with Tukey’s test (p<0.05). Lh, low humidity. Hh, high humidity. **c**, A schematic illustration of examination of humidity effect on SA signaling in plants. **d**, RT-qPCR analysis of BTH-induced *PR1* and *PR2* expression level in the *sid2-2* mutant plant under Lh or Hh, at 0 h and 24 h after 100 μM BTH spray. Plants were pre-treated with Lh or Hh for 24 h before BTH treatment. Data represent mean ± SEM (n = 3 biological replicates). Different letters indicate statistically significant differences, as analyzed by two-way ANOVA with Tukey’s test (p<0.05). **e**, High humidity does not affect the oligomer/monomer ratio of NPR1 protein. The *pNPR1:NPR1-YFP/nprl-6* plants were pre-treated with Lh or Hh for 24 h, sprayed with 100 μM BTH and sampled 12 h after BTH treatment. Proteins were extracted with (+) or without (-) DTT (50 mM) in the sample buffer and subject to SDS-PAGE. NPR1 was detected by immunoblot using monoclonal anti-GFP antibody. Both oligomeric (O) and monomeric (M) forms of NPR1-YFP were detected. Asterisk indicates non-specific band. Actin was used as loading control. **f**, Western blot of NPR1-YFP protein in the total, cytosolic and nuclear fractions from the *pNPR1:NPR1-YFP/npr1-6* plant. Plants were pre-treated with different humidity for 24 h, sprayed with 100 μM BTH, and sampled at different time points after BTH treatment. Anti-GFP antibody was used to detect NPR1-YFP protein in each fraction. Rubisco and Histone3 protein were used as loading controls. **g-h**, Confocal microscopy images (**g**) and quantification (**h**) of NPR1-YFP fluorescence in the *pNPR1:NPR1-YFP/npr1-6* plant under high or low humidity. Plants were treated the same as in **f**, and images were taken 24 h after BTH treatment. Scale bar = 50 μm. The fluorescence intensity and nucleus number on z-stacked images were quantified by Image J software. Data in **h** represent mean ± SEM (n = 8 plants), and analyzed by Student’s t-test (**, p<0.01). Experiments were repeated at least three times with similar results.

## Targeting of SA biosynthesis and signal transduction

Previous studies showed that activation of SA signaling leads to transcriptional induction of *ICS1* and *CBP60g/SARD1*, two master regulators of SA synthesis, and SA production^32,33^, making the pathway a positive-feedback loop. We therefore attempted to dissect the effect of high humidity on SA biosynthesis or signaling, by using SA pathway mutants. To determine whether high humidity affects SA biosynthesis in the absence of SA signaling, we measured synthesis-related gene expression and hormone level in the *npr1-6* mutant, which is severely blocked in SA signaling, under high or low humidity. As shown in Extended Data Fig. 5a-c, the expression of *ICS1* and *CBP60g* genes and the levels of SA, SAG and Pip were still inhibited under high humidity in the *npr1-6* mutant, suggesting that high humidity inhibits SA biosynthesis independently of SA signaling.

Next, we used *sid2-2*, the SA biosynthesis mutant, to examine whether SA signaling induced by exogenous BTH treatment is affected by high humidity. Results showed that BTH-induced expression of downstream genes, including *PR1* and *PR2*, was still dramatically suppressed under high humidity (Fig. 2c-d), indicating that high humidity inhibits SA signal transduction. Collectively, our results suggest that high humidity likely targets both SA synthesis and signaling processes, which together contribute to the strong inhibition of SA responses under high humidity.

## Overaccumulation of NPR1 protein in the nucleus

We next focused to dissect how high humidity inhibits SA signaling. Our RNAseq results showed that key signaling components such as NPR1, NPR3/4 and TGAs^4^ are not transcriptionally regulated by high humidity (Fig. 1g, Extended Data Fig. 6a). We therefore hypothesized that high humidity affects the protein level of signaling components. NPR1 is a SA receptor and transcriptional co-activator, and works together with TGA transcription factors in the nucleus to activate transcription of downstream genes^6–9^. Previous studies showed that NPR1 protein forms oligomers in the cytosol when SA level is low and SA treatment triggers its release and entry into the nucleus to activate gene expression^34,35^. We examined the level of NPR1 protein in the monomer or oligomer form under different humidity in the *pNPR1:NPR1-YFP/npr1-6* plants, and no significant difference was found (Fig. 2e). Similar trends were observed in the *35S:NPR1-YFP/sid2-2* plant, in which the endogenous SA synthesis is almost abolished and humidity regulation of BTH-induced NPR1 protein level can be assessed (Extended Data Fig. 6b). These results suggest that high humidity does not affect the oligomer-to-monomer transition of NPR1 protein.

We then performed fractionation of total protein extracts and detected NPR1 protein in the cytosol or nucleus fraction under different humidity, in the *pNPR1:NPR1-YFP/npr1-6* plants. Intriguingly, NPR1 protein in the nucleus accumulated to a significantly higher level under high humidity, compared to low humidity (Fig. 2f)). Noticeably, the increase of nucleus-located NPR1 protein was more profound at 12 h, compared to 24 h, after BTH treatment. We also detected the cytosolic and nuclear amount of NPR1 protein in the *35S: NPR1-YFP/sid2-2* plants and got similar results (Extended Data Fig. 6c). In addition, we detected the fluorescence signal of NPR1-YFP protein, by confocal microscopy, under different humidity in *pNPR1:NPR1-YFP/npr1-6* plants. We could not detect a clear NPR1-YFP signal 12 h after BTH treatment, likely due to the different detection threshold for western blot vs confocal microscopy. Yet, 24 h after BTH treatment, a stronger fluorescence intensity as well as a higher number of fluorescent nuclei were detected in plants under high humidity (Fig. 2g-h, Extended Data Fig. 6d-e).

This higher NPR1 protein abundance under high humidity was surprising and seemed counterintuitive to the lower SA responses under that condition. Nonetheless, we went on to examine the binding affinity of NPR1 protein, as a transcriptional co-activator, to the target gene promoter under different humidity by chromatin immune-precipitation (ChIP)-qPCR. The nuclei fractions of the *pNPR1:NPR1-YFP/npr1-6* plants under different humidity were isolated for ChIP, and the amount of DNA fragments associated to *PR1* gene promoter and pulled down by NPR1-YFP protein was analyzed. Interestingly, our results indicated a significantly reduced binding of NPR1 protein to different regions of the *PR1* promoter under high humidity, compared to low humidity (Fig. 3a), suggesting that the over-accumulated NPR1 proteins under high humidity are poorly active.

**Fig. 3.**
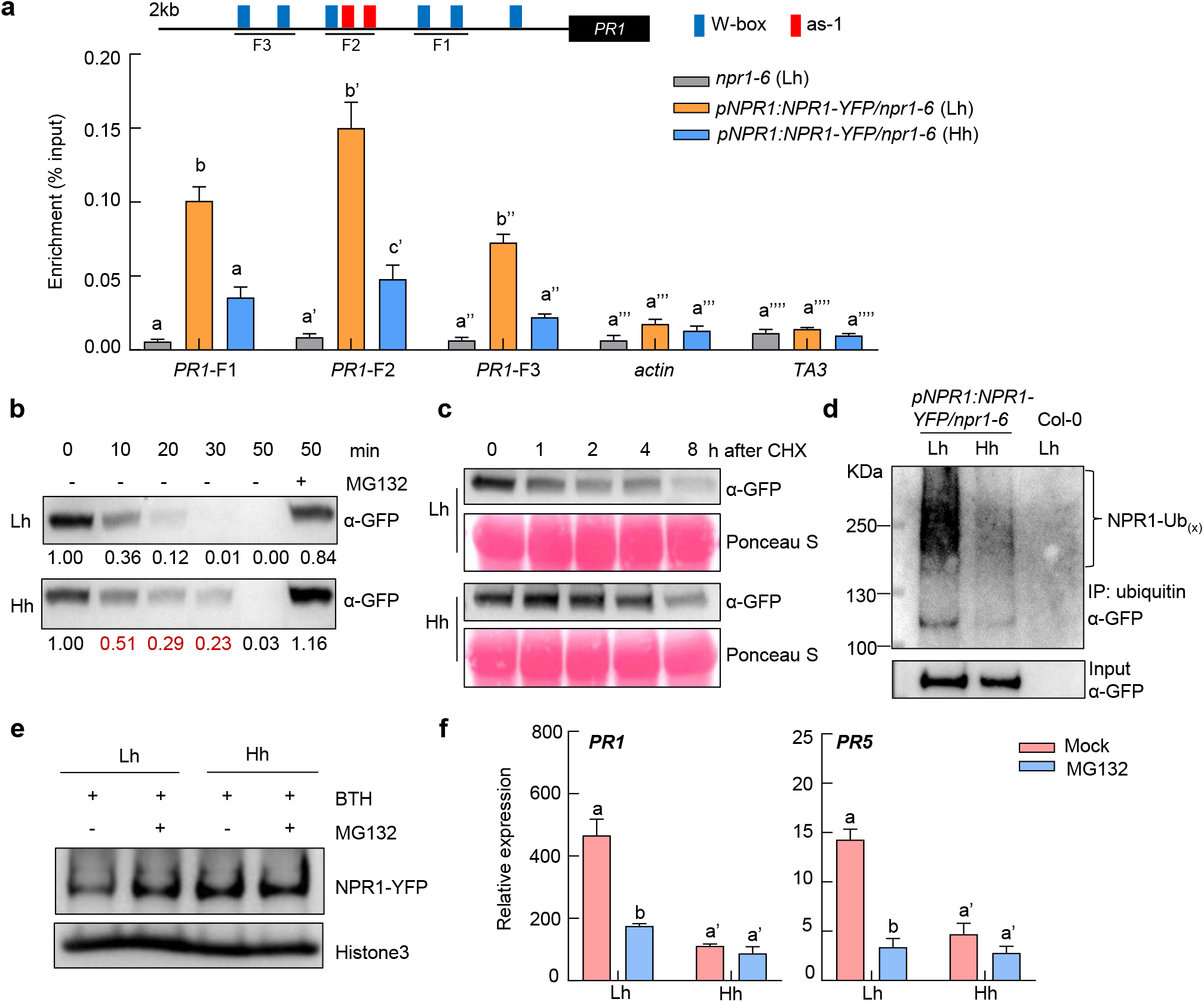
NPR1 protein showed a lower binding affinity to target gene promoter, as well as a reduced level of ubiquitination and proteasome-mediated degradation under high humidity. **a**, NPR1 protein has a lower binding affinity to *PR1* gene promoter under high humidity, as shown by ChIP-qPCR. The *pNPR1:NPR1-YFP/npr1-6* plants were pre-treated with Lh or Hh for 24 h, infiltrated with 50 μM MG132 solution, air-dried for ~0.5 h, and then sprayed with 100 μM BTH. Samples were taken 12 h after BTH treatment. The *npr1-6* plants was used as negative control. ChIP was performed using anti-GFP antibody and qPCR was performed to amplify different regions of *PR1* gene promoter. The *actin* and *TA3* genes were used as negative control. Results from three experimental repeats are shown. Data represent mean ± SEM (n = 3 experimental replicates). Different letters indicate statistically significant differences, as analyzed by two-way ANOVA with Tukey’s test (p<0.05). **b**, *In vitro* cell-free degradation assay. The *pNPR1:NPR1-YFP/npr1-6* plants were pre-treated with Lh or Hh for 24 h, sprayed with 100 μM BTH and sampled 12 h later. Total proteins were extracted and incubated for indicated time, with or without 50 μM MG132. NPR1-YFP protein was detected by western blotting using anti-GFP antibody. Band intensity was quantified by Image J. **c**, NPR1 protein degradation *in vivo*. The *pNPR1:NPR1-YFP/npr1-6* plants were treated the same as in **b**, except that, after BTH treatment for 12 h, plants were dipped with 100 μM cycloheximide (CHX) solution and sampled at the indicated time after CHX treatment. NPR1-YFP protein was detected by western blot using anti-GFP antibody. Rubisco indicates equal loading. **d**, NPR1 protein ubiquitination level under different humidity. The *pNPR1:NPR1-YFP/npr1-6* plants were treated the same as in **a**. Ubiquitinated proteins were enriched using anti-ubiquitin beads, and NPR1-GFP (NPR1-Ubx) in input and IP samples were detected by western blot with anti-GFP antibody. **e-f**, MG132 treatment on plants under low humidity phenocopies high humidity effect. The *pNPR1:NPR1-YFP/npr1-6* plants were treated the same as in **a**. **e**, Nuclear proteins were extracted and subject to SDS-PAGE and NPR1-YFP protein was detected by anti-GFP antibody (with Histone3 as loading control). **f**, The *PR* gene expression was analyzed by qRT-PCR. Data represent mean ± SEM (n = 3 biological replicates). Different letters indicate statistically significant differences, as analyzed by two-way ANOVA with Tukey’s test (p<0.05). Experiments were repeated at least three times with similar results.

## Disruption of NPR1 protein ubiquitination and degradation

Previous studies showed that NPR1 protein undergoes active degradation via 26S proteasome in the nucleus, which tightly regulates NPR1 abundance and immune strength of plants^36^. To determine whether the increase in nuclear NPR1 protein level was caused by abnormal protein degradation under high humidity, we performed the cell-free degradation assay, using protein extracts from the *pNPR1:NPR1-YFP/npr1-6* plants after different humidity treatments. As shown in Fig. 3b, the degradation of NPR1 protein under high humidity was dramatically delayed, compared to that under low humidity (Fig. 3b). We also monitored NPR1 degradation rate in leaves treated with cycloheximide (CHX), a protein synthesis inhibitor. Consistently, a slower degradation rate of NPR1 protein was observed under high humidity compared to low humidity (Fig. 3c).

Next, we tested if the delayed NPR1 protein degradation is due to an altered status of ubiquitination, the primary type of post-translational modifications regulating protein stability^37^. The ubiquitination level of NPR1 protein in the *pNPR1:NPR1-YFP/npr1-6* plants under different humidity was examined and, as shown in Fig. 3d, NPR1-YFP protein displayed a much lower ubiquitination level under high humidity compared to low humidity (Fig. 3d). Again, a similar trend was observed in the *35S:NPR1-YFP/sid2-2* plants (Extended Data Fig. 7a). Therefore, high humidity adversely affects the ubiquitination and degradation processes of NPR1 protein.

To further test if the reduction in NPR1 protein degradation is accountable for the impeded SA responses under high humidity, we treated plants with MG132, a 26S proteasome inhibitor^38^, and examined SA response. Our results showed that MG132 treatment, compared to mock, increased NPR1-YFP protein level under low humidity, but not under high humidity. Furthermore, NPR1 protein level under high humidity is comparable to that under low humidity after MG132 treatment (Fig. 3e), suggesting that the 26S proteasome associated with NPR1 degradation is “already” inhibited under high humidity (without MG132). Importantly, MG132 treatment led to a significant inhibition of SA gene expression (i.e., *PR1* and *PR5*) under low humidity, but did not show an obvious effect under high humidity (Fig. 3f). Similar trends were observed in the *35S:NPR1-YFP/sid2-2* plants (Extended Data Fig. 7b-c). Together, these results suggest that a reduction of proteasome-mediated NPR1 degradation under high humidity leads to an over-accumulation of “inactive” NPR1 proteins in the nucleus and dampened SA signaling. Our results are in agreement with the crucial role of the ubiquitin-proteasome system (UPS)-mediated proteolysis of transcriptional activators in stimulating transcription, as demonstrated in many yeast, animal and plant studies^39–46,57–59^ and, in particular, the importance of NPR1 protein turnover in fully activating SA response genes as reported previously^36^.

## Downregulation of Cullin 3-mediated cellular ubiquitination pathway and 26S proteasome pathway

We then explored the mechanism of high humidity suppression on NPR1 degradation. Previous studies showed that NPR1 protein is phosphorylated at multiple sites, which regulates its ubiquitination and protein stability^47^. A protein-pull down experiment followed by mass spectrometry was carried out to identify NPR1 phosphorylation sites under different humidity. Multiple phosphorylation sites, including Y330, S354 and T373, were detected, but the phosphorylation level seemed similar under high or low humidity (Extended Data Table 1), suggesting that the differential NPR1 degradation under different humidity is unlikely due to altered phosphorylation level at these sites. Nonetheless, we found that high humidity resulted in a global down-regulation of cellular ubiquitination level in Arabidopsis leaf after BTH treatment, as revealed by ubiquitin antibody probing of total protein extracts (Fig. 4a). We then examined the transcript level of E1 ubiquitin-activating genes (*UBAs*, 2 in Arabidopsis genome), E2 ubiquitin-conjugating genes (37 in Arabidopsis genome)^48^ and NPR1-associated E3/E4 ubiquitin ligase components^36,49,50^. Strikingly, *UBA1/2*, the two E1 genes in Arabidopsis, and the E3/E4 *genes-Cullin (CUL)3A/B, HOS15* and *UBE4*, which were shown to mediate NPR1 protein ubiquitination and stability^36,49,50^, were transcriptionally repressed under high humidity (Fig. 4b). While the E2s specifically responsible for NPR1 protein ubiquitination are still unknown, we did not observe a global effect of humidity on E2 family genes (Extended Data Fig. 8). We then examined the transcript level of 26S proteasome machinery genes in our RNAseq. Remarkably, almost all the structural components of 26S proteasome^51^, including the 20S Core Protease (e.g., *PAs* and *PBs*), 19S Regulatory Particle (e.g., *RPNs* and *RPTs*) and accessory factors/chaperones (e.g., *UMP1a* and *ECM29*), were transcriptionally suppressed under high humidity (Fig. 4b, Extended Data Fig. 9). The structural genes of the COP9 Signalosome (CSN), which regulates the activity of cullin subunit of E3 ligases by modification of NEDD8^52^, were not obviously regulated by humidity (Fig. 4b, Extended Data Fig. 9). These results suggest that high humidity attenuates the UPS pathway in the plant cell, likely leading to a global effect on ubiquitination and proteolysis.

**Fig. 4.**
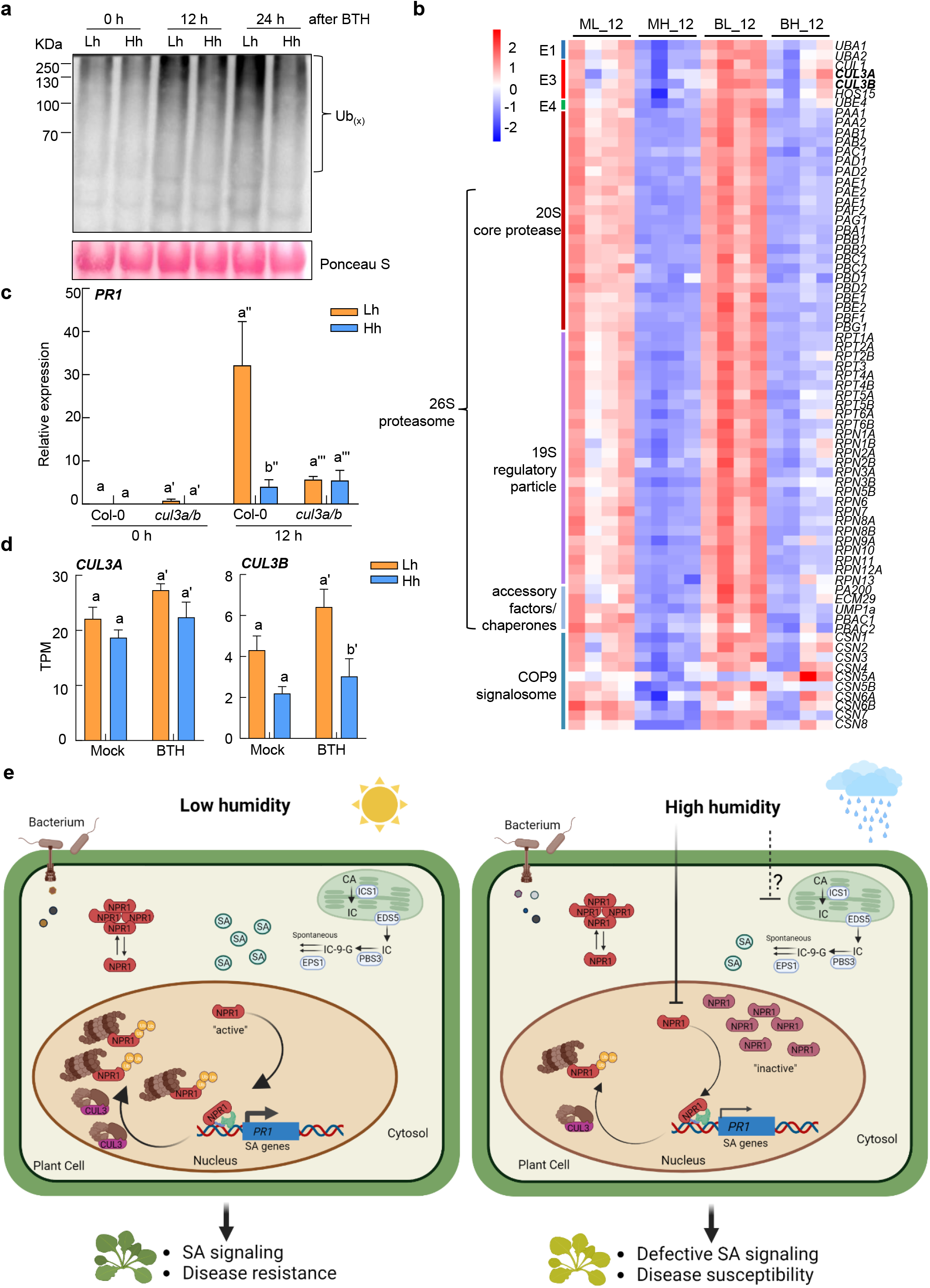
High humidity leads to a down-regulation of Cullin3-based E3 ubiquitin ligase and 26S proteasome pathway in plants. **a**, A global down-regulation of protein poly-ubiquitination under high humidity. Col-0 plants were pre-treated with different humidity for 24 h, sprayed with 100 μM BTH and sampled 12 h or 24 h after BTH. Total proteins were extracted and poly-ubiquitinated proteins were detected by western blot with anti-ubiquitin antibody. **b**, The E1s, NPR1-related E3/E4 and 26S proteasome machinery genes are down-regulated under high humidity, based on our RNAseq analysis. A heatmap based on the TPM of each gene is shown. **c**, RT-qPCR analysis of *PR1* expression level in Col-0 and the *cul3a/b* mutant plants under high or low humidity. Plants were treated the same as in **a** and sampled at 0 h and 12 h after BTH. Data represent mean ± SEM (n = 3 biological replicates). **d**, The *CUL3A* and *CUL3B* expression level under different humidity after mock or BTH treatment in our RNAseq. TPM, transcript per million reads. Data represent mean ± SEM (n = 4 biological replicates). For results in **c** and **d**, different letters indicate statistically significant differences, as analyzed by two-way ANOVA with Tukey’s test (p<0.05). **e**, A working model illustrating findings in this study. Under low humidity (left panel), upon pathogen infection, SA is rapidly produced and perceived by SA receptors, including NPR1. NPR1 protein is ubiquitinated by CUL3 E3 ubiquitin ligase-mediated pathway and targeted to the proteasome for degradation in the nucleus. Clearance of the “exhausted” ubiquitinated NPR1 from the target gene promoter potentially allows fresh proteins to re-initiate transcription, which is important for full SA signaling and disease resistance of plants. Under high humidity (right panel), both SA accumulation and signaling responses are suppressed. The CUL3 and 26S proteasome machinery components are transcriptionally inhibited, leading to impairment in NPR1 ubiquitination and degradation. This results in over-accumulation of “inactive” NPR1 proteins, which failed to bind the target gene promoter effectively, in the nucleus, and consequently defective SA signaling and disease susceptibility of plants. The mechanism by which SA biosynthesis is suppressed under high humidity is still unknown. CA, chorismate. IC, isochorismate. IC-9-G, isochorismate-9-glutamate. ICS1, isochorismate synthase 1. PBS3, *avrPphB* susceptible 3. EPS1, enhanced *Pseudomonas* susceptibility 1.

We next aimed to identify which E3 ubiquitin ligase is biologically relevant to the reduced NPR1 degradation under high humidity. We obtained individual mutants of the E3/E4s known to regulate NPR1 protein ubiquitination, *cul3a/b, hos15* and *ube4*^36,49,50^, and examined NPR1 target gene expression under different humidity. Interestingly, while high humidity still suppressed the *PR1* gene expression in the *hos15* and *ube4* plants, the expression of *PR1* was similar under high or low humidity in the *cul3a/b* mutant plant (Fig. 4c, Extended Data Fig. 10), implying that high humidity suppression of NPR1 target gene is attributed to and relies on CUL3A/B. Consistently, the *CUL3B* transcript was reduced under high humidity in our RNAseq (Fig. 4d). Taken together, these results suggest that a global down-regulation of the cellular ubiquitination pathway, including CUL3-based E3 ubiquitin ligase, and the 26S proteasome pathway compromises NPR1 functionality and SA responses under high humidity (Fig. 4e).

## Discussion

Recent years witnessed tremendous advances in understanding the sensing mechanisms and physiological output of climate factors in regard to different aspects of plant life, including resistance against infectious microbes^53–56^. Our study focuses on air humidity influence on plant immunity, a previously under-studied question, and defines SA pathway and NPR1 protein turnover as biologically relevant targets of high air humidity, disruption of which leads to susceptibility and poor plant health under pathogen attack. Our results also highlight the UPS-mediated proteolysis of transcription co-activator, which potentially eliminates the “spent” proteins and allows fresh proteins to re-initiate transcription as proposed in previous studies across kingdoms^36,39–46,57–59^, as a key process underlying humidity interference with plant resistance.

While SA signaling is strongly inhibited by high humidity, our results suggested that hormone production *per se* is also likely to be suppressed under high humidity, since in the *npr1-6* mutant plant, the level of SA and, more profoundly, Pip is still inhibited under high humidity (Extended Data Fig. 5). It will be interesting to study in the future whether the SA/Pip synthesis steps (e.g., the level of SA precursors) are negatively regulated under high humidity. In addition, whether there are NPR1-independent mechanisms by which high humidity suppresses SA signaling (e.g., through down-regulation of UPS, in light of the importance of an intact ubiquitination pathway in stimulating plant immunity^60^) is also interesting to explore. Furthermore, signaling events upstream of high humidity suppression of SA pathway, namely the humidity sensing mechanisms, are completely unknown at the moment and should be important to study. In-depth understanding of the environmental influences on plants should assist in the development of novel approaches of safeguarding plant immune capacity and better coping plants with adverse climate conditions.

## Methods

### Plant materials, growth condition, and humidity treatment

All Arabidopsis plants used in this study were in the Columbia-0 genetic background. Seeds were surface-sterilized with 5% sodium hypochlorite solution for 10 min and then washed with sterile water. Seeds were cold-treated for 2 days before sowing on soil pots. All plants were grown in a growth chamber (Hettich) with the temperature setting of 22°C, 12 h light/12 h dark photoperiod and 60% relative humidity. Plants were grown to 3.5-4 weeks for experiments.

The *npr1-6* (SAIL_708F09)^31^, *sid2-2*^30^, *hos15*^49^. *ube4*^50^ and *uba1*^60^ mutants were previously characterized and reported. The *cul3a/b* mutant was generated by crossing of a *cul3a* knockout allele (SALK_050756)^61^ with a *cul3b* knockdown allele (SALK_098014)^36^. To generate the *pNPR1:NPR1-YFP/npr1-6* plant, the *pNPR1:NPR1-YFP* construct^31^ was mobilized into *Agrobacterium tumefaciens* GV3101 strain and transformed into the *npr1-6* plant by floral dip. To generate the *35S:NPR1-YFP/sid2-2* plants, the coding sequence of the *NPR1* (AT1G64280) was amplified from cDNA derived from Col-0 plant and cloned into the gateway entry vector pENTR/D-TOPO (Invitrogen) and then transferred into the destination vector pGWB605 vector^62^. The binary vector was mobilized to *Agro. tumefaciens* GV3101 strain, and the *35S:NPR1-YFP* expression cassette was transformed into the *sid2-2* mutant plant by floral dip method.

For humidity treatment, plants were grown under ~60% RH till 3.5-4 weeks old and then placed in two chambers (MMM, Germany), in which humidity was set to 45-50% RH (for low humidity) or 95% RH (for high humidity). Temperature was set at 22 °C and photoperiod at 12 h light/12 h dark in both chambers. LED lights were used as light source in these chambers.

### Chemical treatment

For experiments with elicitor treatment, plants were pre-treated with different humidity for 24 h before being sprayed with the following chemical: BTH (Sigma; 100 μM, dissolved in 0.1% DMSO supplemented with 0.01% Silwet L-77), MeJA (Sigma; 100 μM, dissolved in 0.1% EthOH supplemented with 0.01% Tween20) or flg22 (Phyto Tech; 200 nM, dissolved in ddH_2_O supplemented with 0.01% Silwet L-77). Treated plants were placed back to different humidity setting and leaf tissue was sampled at different time points for protein detection, transcript measurement, cellular fractionation, confocal imaging, hormone quantification or other assays.

### Bacterial disease assay

*Pst* DC3000 bacteria were grown in LM liquid medium supplemented with 50 mg/L Rifampicin for overnight. The next morning, bacteria were collected by centrifugation at 2500 x *g* for 5 min, washed with sterile water and resuspended. The concertation was adjusted to 1×10^6^ cfu/mL (OD=0.002). Bacteria were infiltrated into Arabidopsis leaves with a needleless syringe, and the infiltrated plants were kept under ambient humidity for about 1 h for water to evaporate and then put into the chamber with controlled humidity for disease development. Disease symptom and bacteria population were recorded and quantified three days later.

### RNA extraction, cDNA synthesis, and RT-qPCR

Leaf samples were snap-frozen by liquid nitrogen, and grounded powders were homogenized in the Trizol reagent (Invitrogen). Total RNA was extracted according to the manufacture’s instructions. RNA samples were quantified using a NanoDrop spectrophotometer (Thermo Scientific). Reverse transcription was performed using the ReverTra Ace qPCR RT Master Mix with gDNA remover (TOYOBO), and qPCR was performed using the SYBR Green Realtime PCR Master Mix (TOYOBO) on a CFX real-time machine (Bio-Rad). The *PP2AA3* gene was used as the reference gene for normalization. All primers for qPCR are listed in Extended Data Table 2.

### RNA-Sequencing and data analysis

For RNA-seq experiments, Col-0 plants were treated with high or low humidity for 24 h, sprayed with 100 μM BTH and placed back to different humidity setting. Leaves were collected 12 h after BTH/Mock treatment and 4 leaves were collected as one biological replicate. Four biological replicates were collected for each treatment. Total RNA was extracted using Trizol Reagent (Invitrogen) based on the manufacture’s instructions, and genomic DNA was removed using the DNaseI (Invitrogen). RNA was further purified using the RNeasy MinElute Cleanup kit (QIAGEN). RNA quality check, library construction, and sequencing were performed by the Majorbio company. Briefly, RNA quality was determined by 2100 Bioanalyser (Agilent) and quantified using the ND-2000 (NanoDrop Technologies). RNA-seq library was prepared using the TruSeq TM RNA sample preparation Kit from Illumina (San Diego, CA), using 1 μg of total RNA for each sample. The messenger RNA was isolated according to the polyA selection method by oligo (dT) beads and then fragmented in fragmentation buffer. Double-stranded cDNA was synthesized using a SuperScript double-stranded cDNA synthesis kit (Invitrogen, CA) with random hexamer primers (Illumina). Then the synthesized cDNA was subjected to end-repair, phosphorylation, and ‘A’ base addition according to Illumina’s library construction protocol. Libraries were size selected for cDNA target fragments of 300 bp on 2% Low Range Ultra Agarose followed by PCR amplification using Phusion DNA polymerase (NEB) for 15 cycles. After quantified by TBS380 (Picogreen), the paired-end sequencing library was sequenced with the Illumina NovaSeq 6000 sequencer (2 × 150bp read length).

For data analysis, raw paired-end reads were trimmed and quality controlled by Fastp with default parameters^63^. Then clean reads were separately aligned to reference genome with orientation mode using HISAT2 software^64^. The mapped reads of each sample were assembled by StringTie in a reference-based approach^65^. The expression level of each transcript was calculated according to the transcripts per million reads (TPM) method, and RSEM was used to quantify gene abundance^66^. The differential expression analysis was performed by the DEGSeq2 with default parameters^67^.

### Phytohormone extraction and quantification

Phytohormone extraction and quantification were performed based on previous reports with slight modification^31^. Leaf tissue was frozen and ground in liquid nitrogen and hormones were extracted at 4°C in the ice-cold extraction buffer (80% methanol in water, 0.1% formic acid, 0.1 g/L butylated hydroxytoluene and 20 nM ABA-d6). The extraction step was repeat twice and total supernatant was speed-dried in a vacuum centrifugal concentrator (Beijing JM Technology). The pellet was resuspended in 30% methanol solution. Hormone levels were quantified using the AB SCIEX 4000Q TARP LC/MS/MS system (SCIEX QTRAP 6500+). Selected ion monitoring (SIM) was conducted in the negative ES channel for SA (137.0>93.0), SAG (299.0>1377.0), JA (209.0>59.0), JA-Ile (322.1>130.1) and the internal standards Propyl 4-hydroxybenzoate (179.0>137.0), H2-JA (211.1>59.0). Pip (130.0>84.0) and its internal standard L-Valine (118.0>72.0) were conducted in the positive ES channel. Parent>daughter SIM pairs, the optimal source cone and collision energy voltages for each molecular were determined by the QTRAP 6500+. All hormone concentrations were normalized by sample fresh weight (FW) in g.

### Nuclear fractionation and western blotting

Plant nuclei were isolated from plants using the CelLytic PN Isolation/Extraction Kit (Sigma) and nine leaf discs (0.75 cm in diameter) were collected per sample/treatment. And 100 μM MG132 (Selleck) and 1 x EDTA-free protease inhibitor cocktail (Roche) were freshly added in the extraction buffer. Following protein extraction and fractionation, total and cytosolic fractions were mixed with 4 x LDS sample buffer and nuclear fraction was resuspended in 60 μL 1 x LDS sample buffer. The α-GFP antibody (Abmart) was used to detect the NPR1-YFP protein and α-Histone 3 antibody (Agrisera) was used to detect Histone 3 (as loading control), and Rubisco was detected by Ponceau S staining.

### Confocal laser scanning microscopy

Confocal images were taken by a Leica TCS SP8 STED system with Z stack program. YFP was excited with an argon laser using a 488 nm beam splitter, and emission was detected with a 520-580 nm bandpass filter. All confocal images were taken using the same paraments. Fluorescence quantification was performed on the ImageJ software. Fluorescence intensity on each image was determined with a 594 minimum threshold and the number of nuclei with detectable fluorescence was counted with a > 3-Infinity size setting.

### *In vitro* cell-free degradation assay

Cell-free degradation assays were performed as described previously^36^. Briefly, the *pNPR1::NPR1-YFP/npr1-6* and *35S::NPR1-YFP/sid2-2* plants were pre-treated with different humidity for 24 h and then sprayed with 100 μM BTH, and 36 leaf discs (0.75 cm in diameter) were sampled 12 h later. Leaf tissues were ground in liquid nitrogen and total proteins were extracted in the extraction buffer (25 mM Tris-HCl, pH 7.5, 10 mM MgCl2, 10 mM NaCl, 10 mM ATP, 0.2% TritonX-100, 0.2% CA630 and 5 mM DTT). The extracted proteins were then incubated at room temperature for 0-60 min, in the presence or absence of 100 μM MG132, and samples were taken at different time points. The NPR1-YFP protein was detected by western blot using the GFP antibody. Band intensity was quantified using ImageJ software.

### Detection of NPR1 ubiquitination

Four-week-old plants were pre-treated with different humidity for 24 h and then 100 μM MG132 solution was infiltrated into the leaves. Leaves were air-dried for about 0.5 h and then 100 μM BTH solution was sprayed on both sides of the leaf. Plants were then placed back to different humidity and samples were taken 12 h later. About sixty leaf discs were sampled for each sample. This assay was performed according to pervious report^68^ with some modifications. Total proteins were extracted in extraction buffer, which contains 50 mM Tris-HCl pH7.5, 100 mM NaCl, 10% glycerol, 6 M Urea, 50 μM MG132, 50 μM PR-619 (Abcam), 10 mM Iodoacetamide (Sigma), 1x Protease inhibitor cocktail (Roche). Protein extracts were centrifuged at 12,000 x *g* at 4°C for 5 min and supernatant was taken. Anti-ubiquitin beads (Cell Signaling Technology) were added and incubated for 2 h at 4°C. The beads were washed 3 times with wash buffer (50 mM Tris-HCl pH7.5, 100 mM NaCl, 10% glycerol, 50 μM MG132, 50 μM PR-619, 10 mM Iodoacetamide, 1x Protease inhibitor cocktail), and proteins were eluted in 60 μL of 1x LDS elution buffer. NPR1-GFP protein was detected using the anti-GFP antibody (Abmart).

### ChIP-qPCR analysis

The ChIP experiment was performed as previously reported^53^, with some modifications. The treated plant leaves were collected and fixed in 1% formaldehyde by vacuum infiltration on ice. Then the fixation solution was replaced with 125 mM glycine solution and vacuumed for 5 min. Leaves were washed with ice water three times and flash-frozen in liquid nitrogen, and grounded into powder. About 1.5 g powder per sample were used to extract the pure nuclei according to the previous report^69^. The 1x protease inhibitor cocktail and 50 μM MG132 were added throughout extraction process to increase protein stability. Pure nuclei pellet was suspended in 150 μL of nuclei lysis buffer (50 mM Tris pH 8.0, 10 mM EDTA pH 8.0, 1% SDS, 1x protease inhibitor cocktail, 50 μM MG132) and incubated on ice for 30 min. Then 1250 μL of ChIP dilution buffer (16.7 mM Tris pH 8.0, 167 mM NaCl, 1.2 mM EDTA, 0.01% SDS, 1x protease inhibitor cocktail, 50 μM MG132) were added and the samples were sonicated for 10 min on Bioruptor (Diagenode) at 4°C. Then 400 μL of ChIP dilution buffer and 200 μl of 10% Triton X-100 were added and samples were centrifuged at 12000 x *g* for 15 min to remove debris. For pre-clearing, samples were incubated with 25 μl of protein A beads (Millipore) for 2 h in cold room (100 μL were collected as input). To capture the DNA–protein complex, GFP-Trap magnetic beads (Chromotek) were added and incubated overnight at 4°C. After washing of beads, DNA-protein complexes were eluted with 540 μL elution buffer (50 mM Tris pH 8.0, 10 mM EDTA pH 8.0, 1% SDS) and boiled at 65°C for 30 min. Then 20 μL of 5 M NaCl were added and samples were incubated at 65°C overnight to remove crosslinking. DNA were purified and quantified by qPCR. Primers are listed in Extended Data Table 2.

### Detection of NPR1 phosphorylation sites

The *pNPR1::NPR1-YFP/npr1-6* plants were pre-treated with high or low humidity for 24 h, sprayed with BTH and sampled 12 h later. Total proteins were extracted in 16 mL of extraction buffer (50 mM Tris pH 7.5, 150 mM NaCl, 0.5 mM EDTA, 1 mM DTT, 50 μM NEM, 5% glycerol, 10 mM IAA (Iodoacetamide, Sigma), 5 μM MG132, 25 mM NaF, 1 mM Na2MO4, 1 x protease inhibitor cocktail, 1x phosphatase inhibitor cocktail, 0.5% TritonX-100, 0.5% CA630 and 0.5% SDS) on ice for 30 min. After centrifugation at 14000 x g at 4°C for 10 min, the supernatant lysate was filtered by two layers of Miracloth (Millipore) and diluted with the same volume of extraction buffer without detergent. NPR1-YFP protein was pulled down by anti-GFP beads (Smart-Lifesciences) with agitation for 3 h at 4°C. Beads were washed three times with extraction buffer without detergent. Eluted proteins were separated on a 10% polyacrylamide gel and stained with the Fast Silver Stain Kit (Beyotime Biotech). The NPR1-YFP band was cut and in-gel digested by trypsin (Promega). The phosphorylated peptides of NPR1 protein were detected by LC-MS/MS at the Mass Spectrometry facility at CAS for Excellence in Molecular Plant Sciences.

### Data availability

The RNA-seq data have been deposited into the NCBI Gene Expression Omnibus under accession GSE210893. All other data are available in the main text or supporting materials.

## Acknowledgments

We would like to thank all the members in Xin lab for helpful discussions. We thank the Greenhouse, Confocal Microscopy Imaging facility and Metabolomics-Mass Spectrometry facility at the CAS Center for Excellence in Molecular Plant Sciences for support in plant growth, confocal imaging and hormone quantification. The *pNPR1:NPR1-YFP* construct was kindly provided by Dr. Sheng Yang He’s lab at Duke University, USA. This research was supported by Chinese Academy of Sciences, Center for Excellence in Molecular Plant Sciences/Institute of Plant Physiology and Ecology, National Key Laboratory of Molecular Plant Genetics, Shanghai Pilot Program for Basic Research – Chinese Academy of Science, Shanghai Branch (JCYJ-SHFY-2021-007) and Chinese Academy of Sciences Strategic Priority Research Program (Type-B; project number: XDB27040211). Lingya Yao is supported by the Youth Program of National Natural Science Foundation of China (NSFC) (project number: 32100238).

## Author contributions

L.Y., Z.J., and X-F.X. conceptualized the project. L.Y. and Z.J. performed most of the experiments, including RNAseq, disease assay, RT-qPCR, protein detection, hormone quantification, confocal imaging and ChIP-qPCR. Y.W. helped with the ChIP-qPCR experiment. S.W. performed JA gene expression. L.Y., Z.J. and X-F.X. wrote the manuscript with input from all authors.

## Competing interests

The authors declare no competing interests.

## Materials & correspondence

Correspondence and material requests should be addressed to.

---

**Extended Data Table 1.**
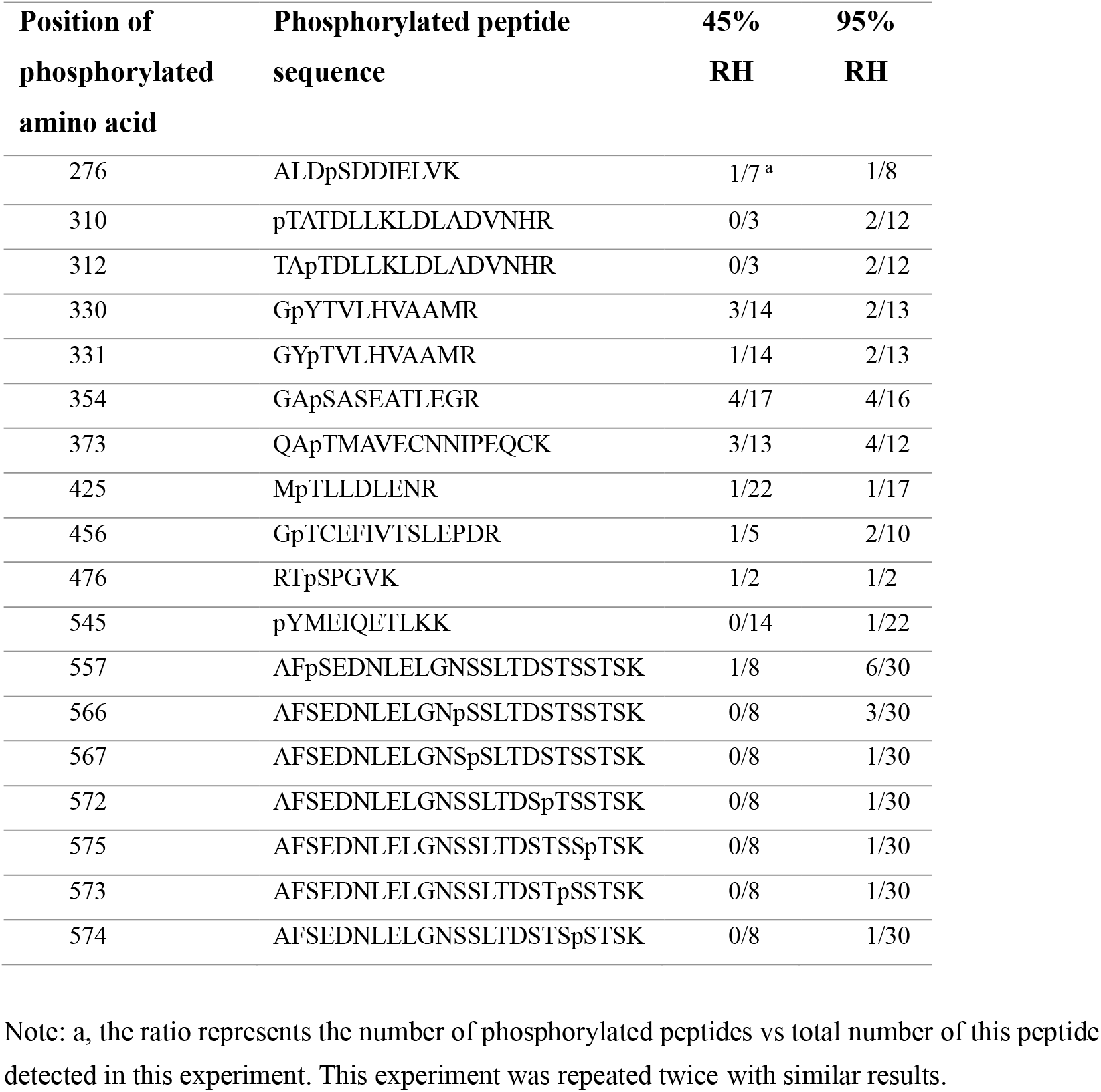
Phosphorylation sites of NPR1 protein under different humidity, as detected by Mass Spectrometry.

**Extended Data Table 2.**
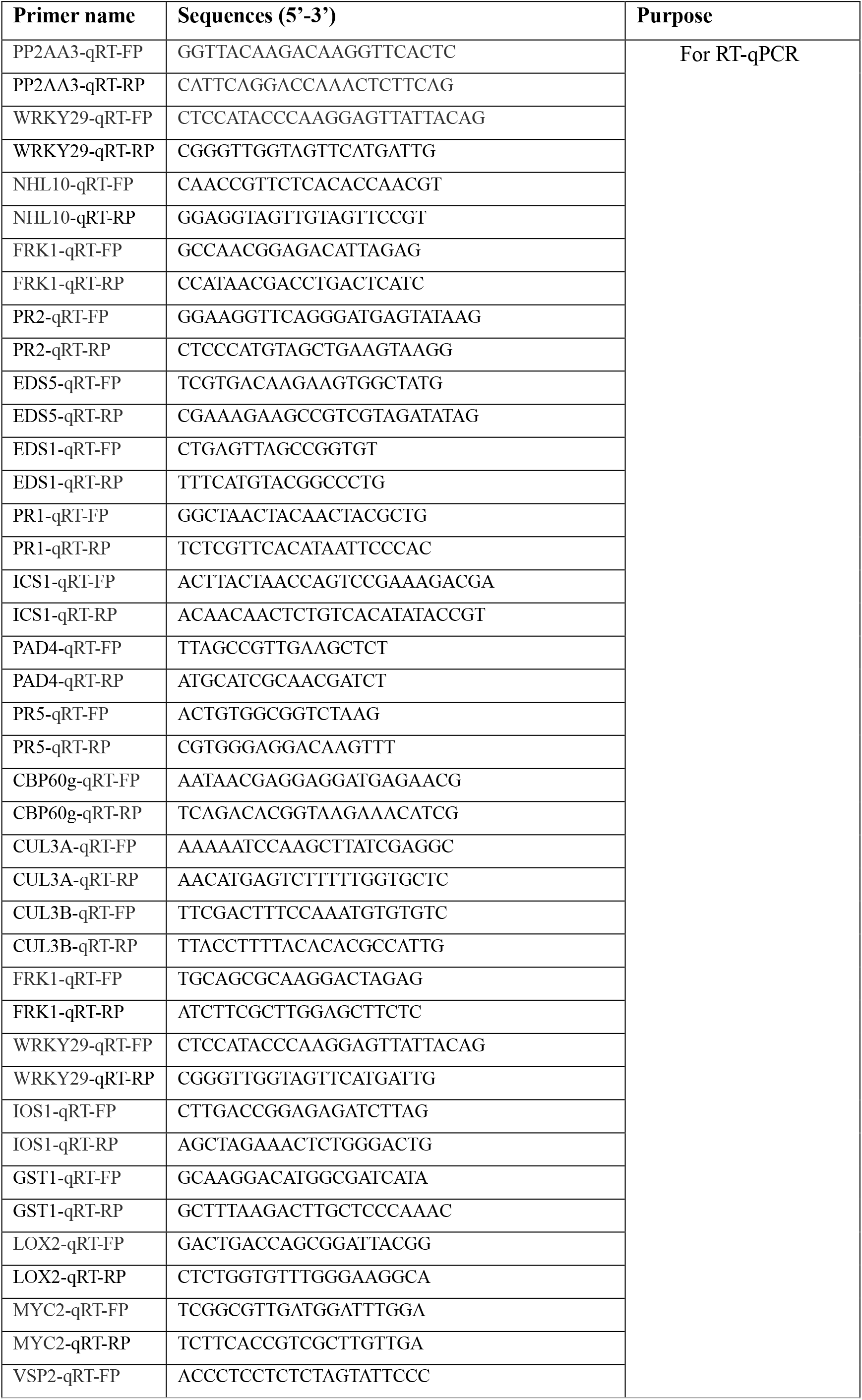
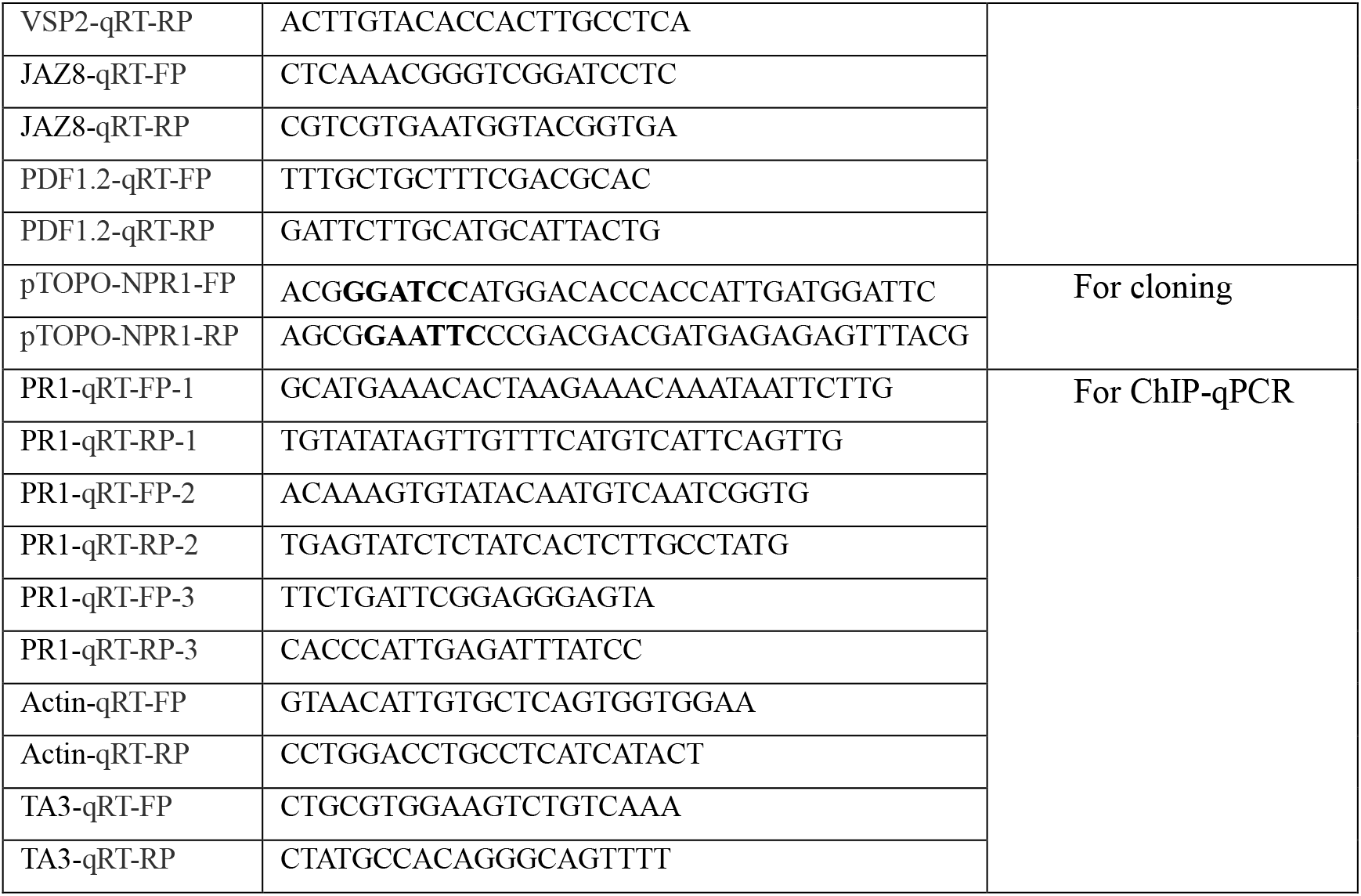
Primers used in this study.

**Extended Data Fig. 1.**
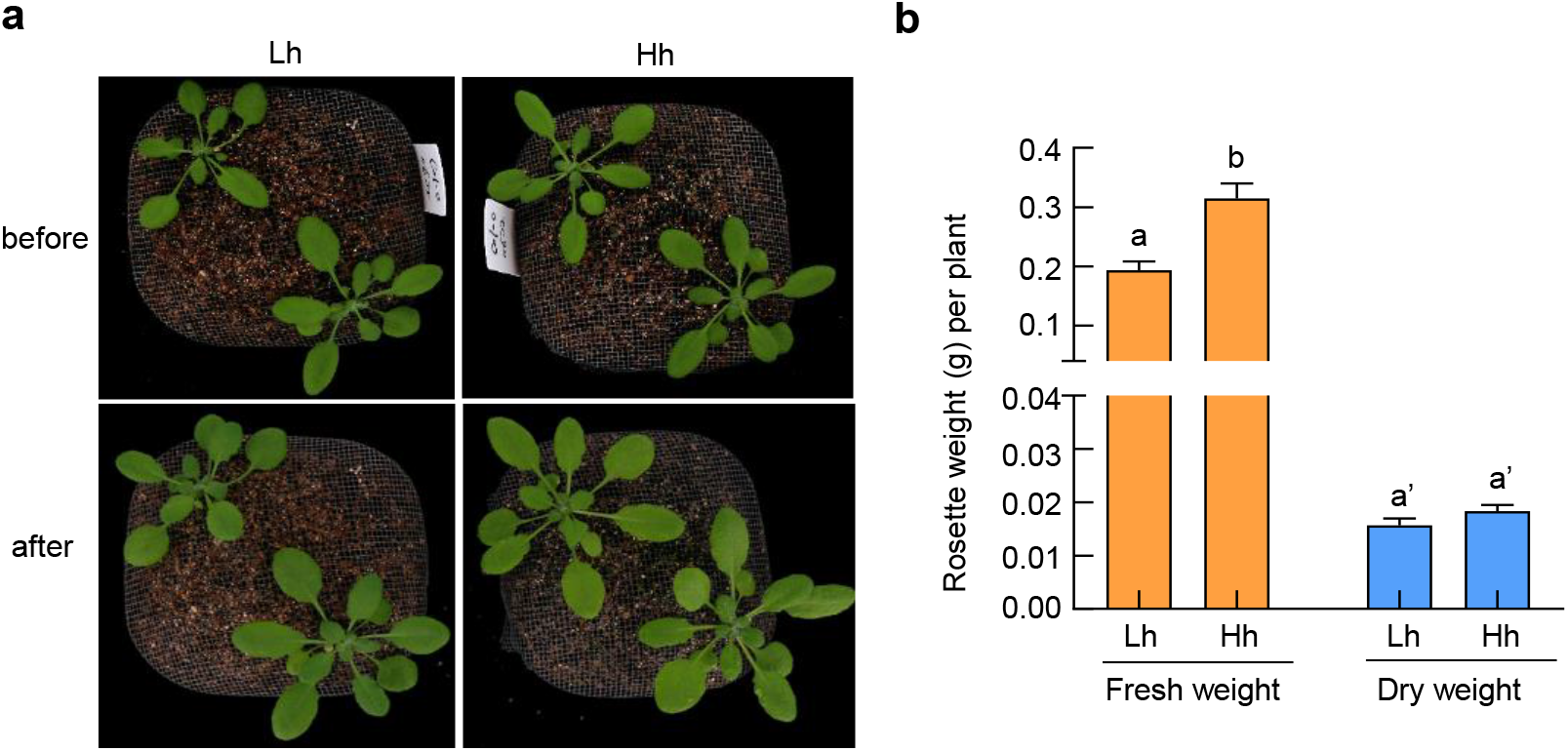
Growth and morphology phenotype of plants under different humidity. **a**, Three-to-four-week old Col-0 plants were treated with low humidity (Lh, 45% RH) or high humidity (Hh, 95% RH) for 2 days. Pictures were taken before and after humidity treatment. **b**, Leaf weight of plants after different humidity treatment. Data are shown as mean ± SEM (n = 4 plants). Different letters indicate statistically significant differences, as analyzed by two-way ANOVA with Tukey’s test (p<0.05).

**Extended Data Fig. 2.**
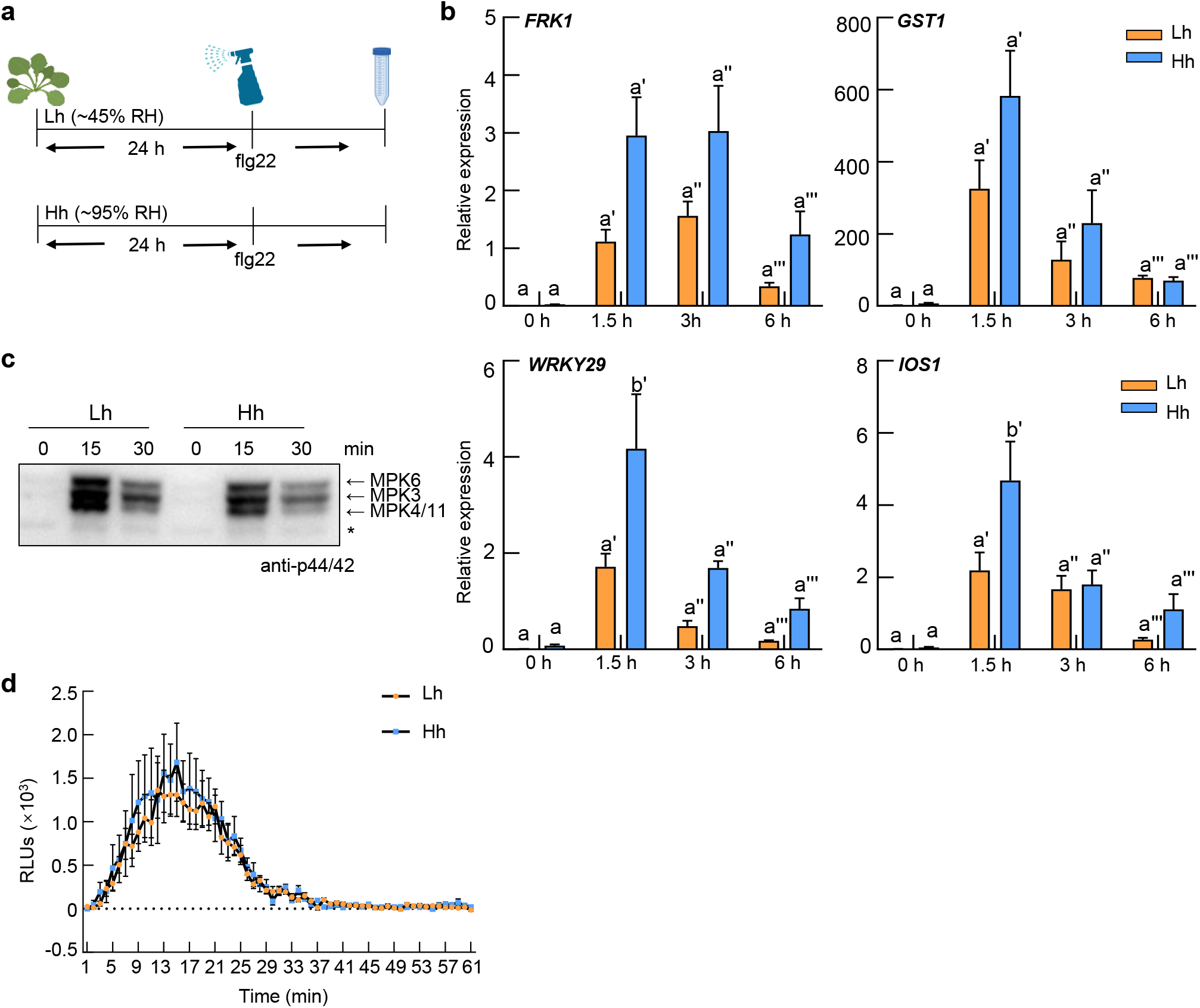
flg22-induced PTI responses were not obviously affected by high humidity. **a**, Schematic diagram showing humidity and flg22 treatment for experiments in this figure. **b**, RT-qPCR analysis of 200 nM flg22-induced *FRK1, GST1, WRKY29* and *IOS1* expression level in Col-0 plants under Lh or Hh. Samples were taken at 0 h, 1.5 h, 3 h and 6 h after flg22 treatment. Data represent mean ± SEM (n = 3 biological replicates). Different letters indicate statistically significant differences, as analyzed by two-way ANOVA with Tukey’s test (p<0.05). **c**, flg22-induced MAPK Phosphorylation in Col-0 plants under Lh or Hh. An equal amount of total protein was loaded in each lane. Asterisk indicates non-specific band. **d**, flg22-induced ROS production in Col-0 plants under different humidity. Data are shown as mean ± SEM (n = 6 leaf discs). Experiments were repeated at least three times with similar trends.

**Extended Data Fig. 3.**
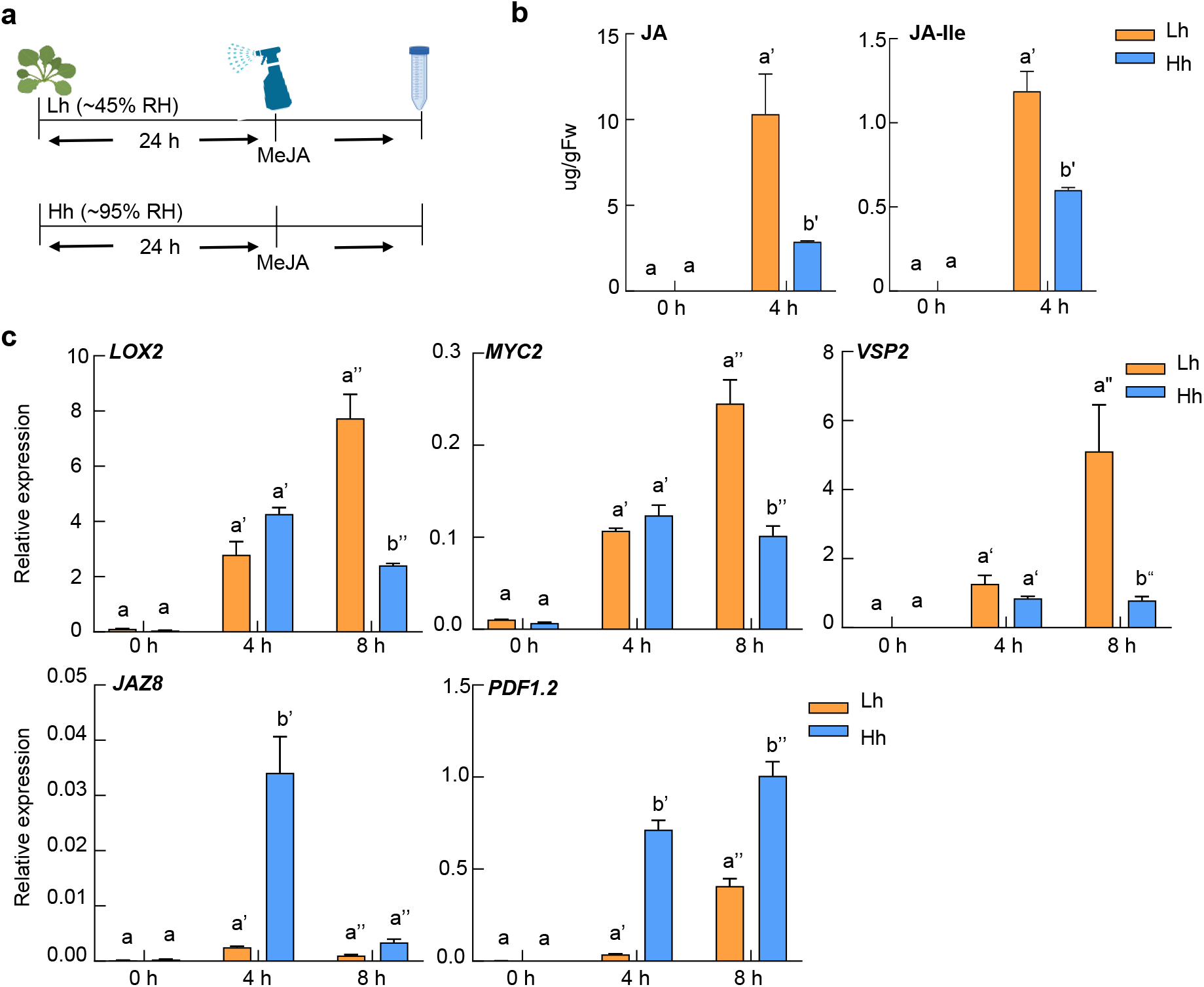
High humidity shows an inconsistent effect on JA responses. **a**, Schematic diagram showing humidity and MeJA treatment for experiments in this figure. **b**, Quantification of JA and JA-Ile in Col-0 plants under Lh or Hh, at 0 h and 4 h after 100 μM MeJA treatment. Data represent mean ± SEM (n = 3 biological replicates). Different letters indicate statistically significant differences, as analyzed by two-way ANOVA with Tukey’s test (p<0.05). **c**, RT-qPCR analysis of JA-responsive gene expression in Col-0 plants under Lh or Hh, after 100 μM MeJA treatment. Data represent mean ± SEM (n = 3 biological replicates). Different letters indicate statistically significant differences, as analyzed by twoway ANOVA with Tukey’s test (p<0.05). Experiments were repeated three times with similar results.

**Extended Data Fig. 4.**
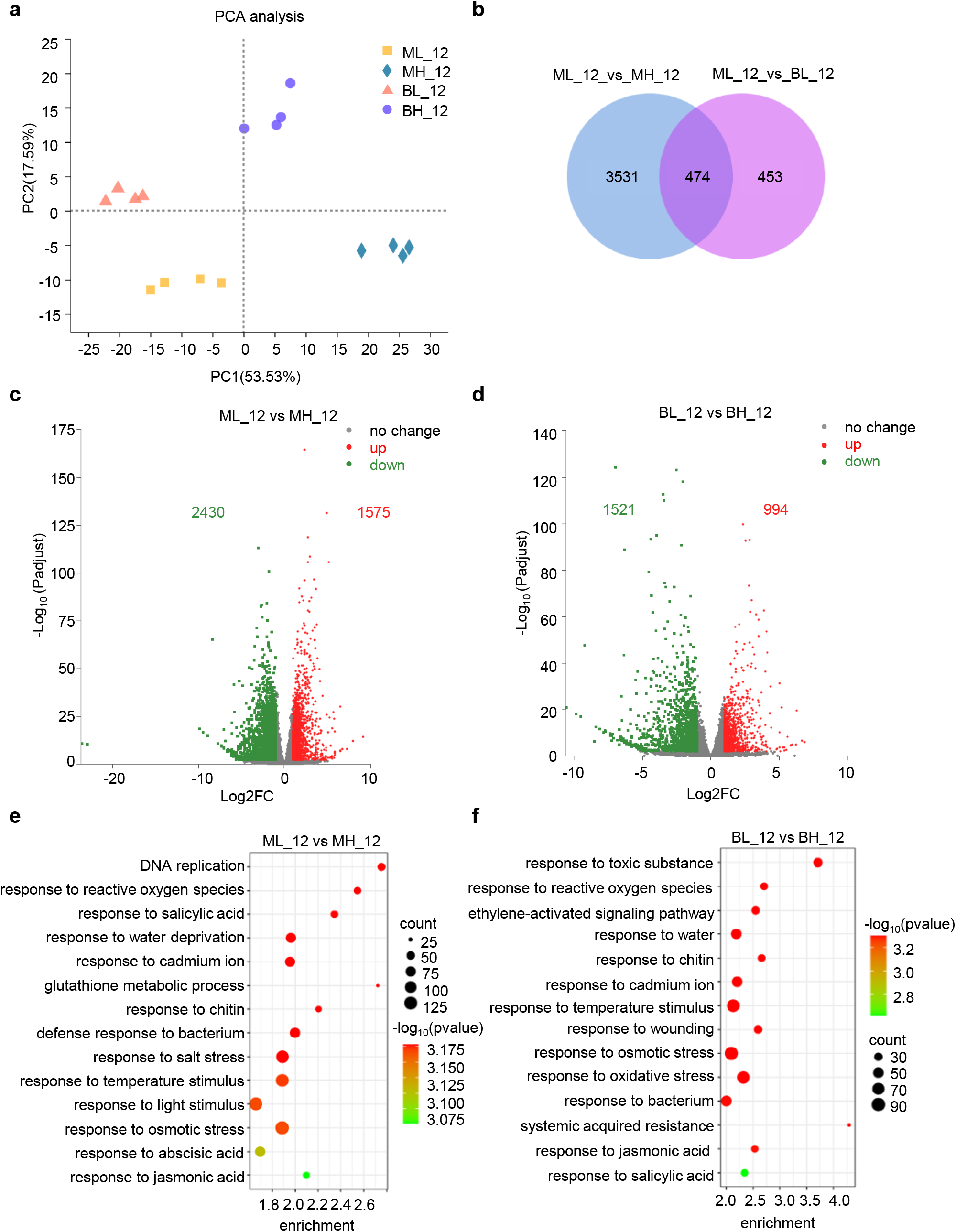
Genome-wide expression analysis of Arabidopsis Col-0 plants under different humidity and BTH treatment. **a**, Principal component analysis. **b**, Venn diagram illustrating the number of genes differentially regulated under Lh or Hh. **c-d**, Volcano map of differentially regulated genes under high and low humidity. **e-f**, GO enrichment analysis of the differentially regulated gene in **c** and **d**. ML_12, 12 h after mock treatment under low humidity. MH_12, 12 h after mock treatment under high humidity; BL_12, 12 h after BTH treatment under low humidity; BH_12, 12 h after BTH treatment under high humidity.

**Extended Data Fig. 5.**
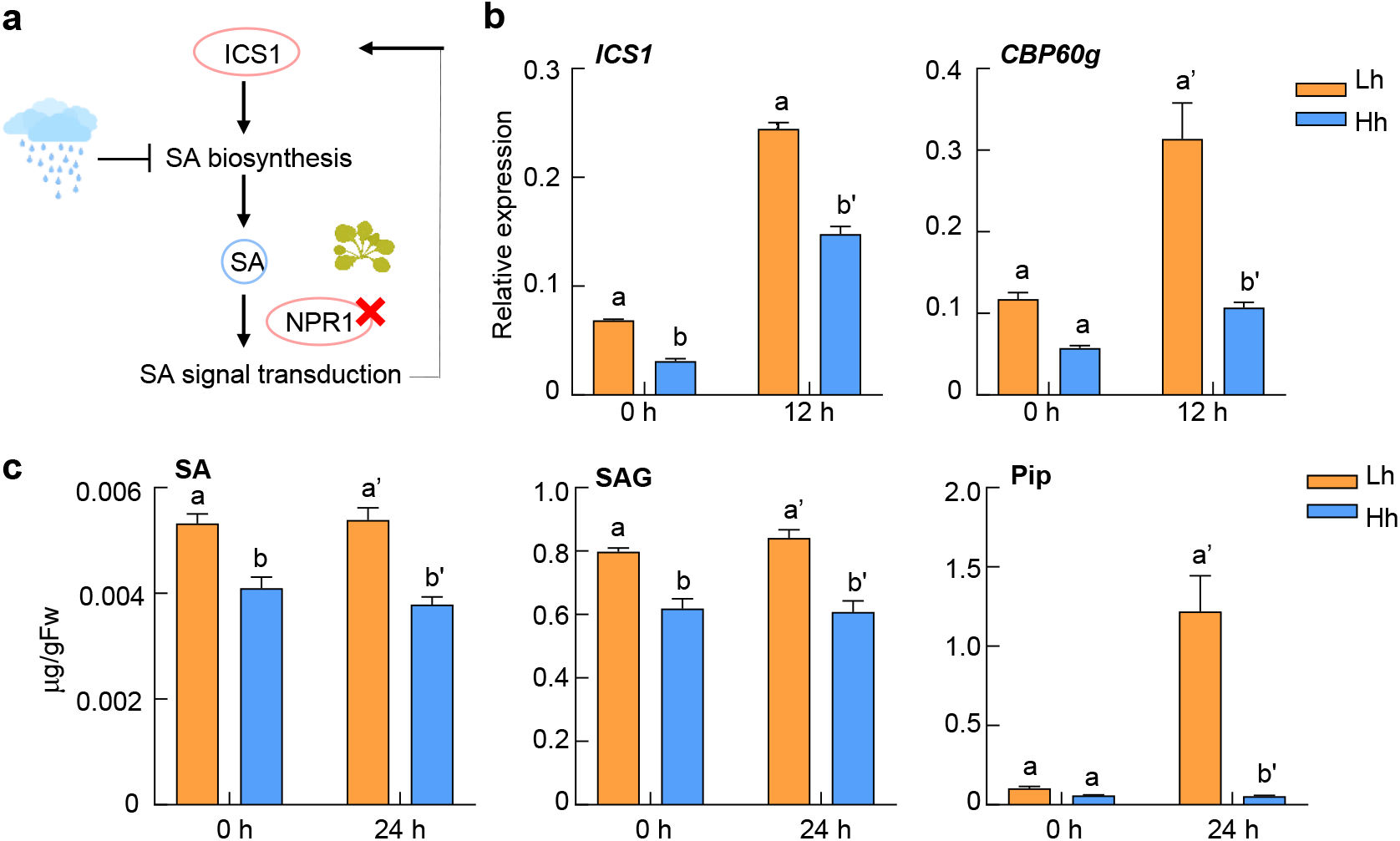
SA biosynthesis is inhibited in the *npr1-6* mutant plant under high humidity. **a**, Schematic diagram of examination of humidity effect on SA biosynthesis. **b**, RT-qPCR analysis of *ICS1* and *CBP60g* expression level in the *npr1-6* mutant under different humidity, at 0 h and 12 h after 100 μM BTH treatment. Plants were pre-treated with Lh or Hh for 24 h. Data were mean ± SEM (n = 3 biological replicates). Different letters indicate statistically significant differences, as analyzed by two-way ANOVA with Tukey’s test (p<0.05). **c**, The SA, SAG and Pip levels in the *npr1-6* mutant under different humidity. Plants were treated the same as in **b**. Data represent mean ± SEM (n = 4 biological replicates). Different letters indicate statistically significant differences, as analyzed by twoway ANOVA with Tukey’s test (p<0.05). Experiments were repeated three times with similar results.

**Extended Data Fig. 6.**
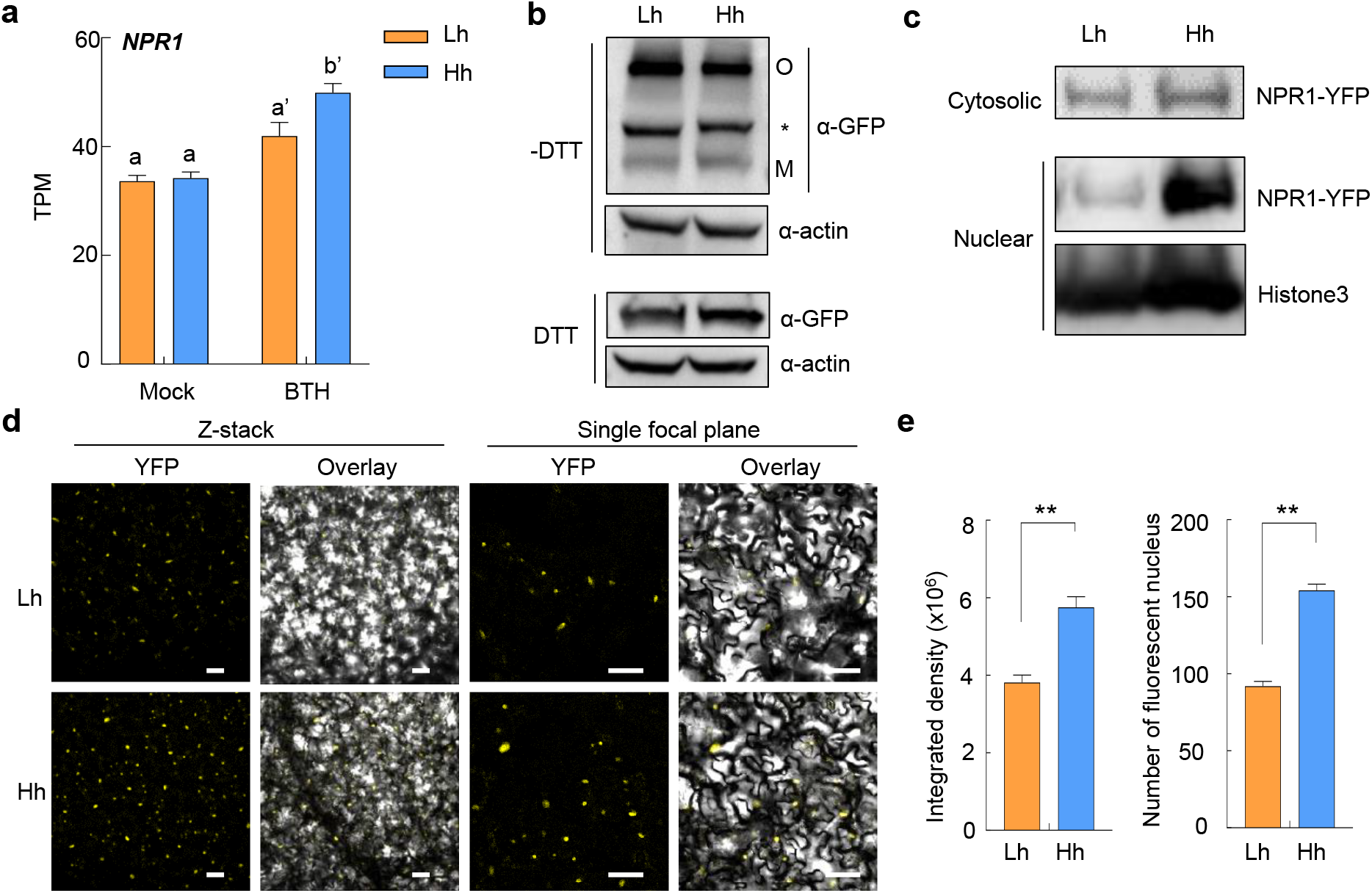
High humidity promotes NPR1 protein accumulation in the nucleus of the *35S:NPR1-YFP/sid2-2* plants. **a**, *NPR1* gene expression is not dramatically changed under different humidity, based on our RNAseq results. TPM, transcript per million reads. Data represent mean ± SEM (n = 4 biological replicates). Different letters indicate statistically significant differences, as analyzed by two-way ANOVA with Tukey’s test (p<0.05). **b**, The oligomer-to-monomer transition of NPR1 was not affected at elevated humidity. The *35S:NPR1-YFP/sid2-2* plants were pre-treated with Lh or Hh for 24 h, treated with 100 μM BTH and sampled 12 h after BTH treatment. Total proteins were extracted in the sample buffer with (+) or without (-) DTT (50 mM), and subject to SDS-PAGE. NPR1 protein was detected by immunoblot using a monoclonal anti-GFP antibody. Both oligomeric (O) and monomeric (M) forms of NPR1-YFP were detected. Asterisk indicated the nonspecific protein. Actin was used as a loading control. **c**, Western blot of NPR1 protein in the cytosolic and nuclear fractions from the *35S:NPR1-YFP/sid2-2* plants. Plants were treated the same as in **b**. Anti-GFP antibody was used to detect NPR1-YFP protein in each fraction, and Histone3 protein was used as loading control. **d-e**, Confocal microscopy images (**d**) and quantification (**e**) of NPR1-YFP fluorescence in the *35S:NPR1-YFP/sid2-2* plant under Lh or Hh. Plants were treated the same as in **b**. Images of YFP channel and bright field were taken 24 h after BTH treatment. Scale bar = 50 μm. The fluorescence intensity and nucleus number on z-stacked images were quantified by Image J software in **e**. Data represent mean ± SEM (n = 12 plants) and analyzed by Student’s t-test (**, p<0.01). Experiments were repeated at least three times with similar results.

**Extended Data Fig. 7.**
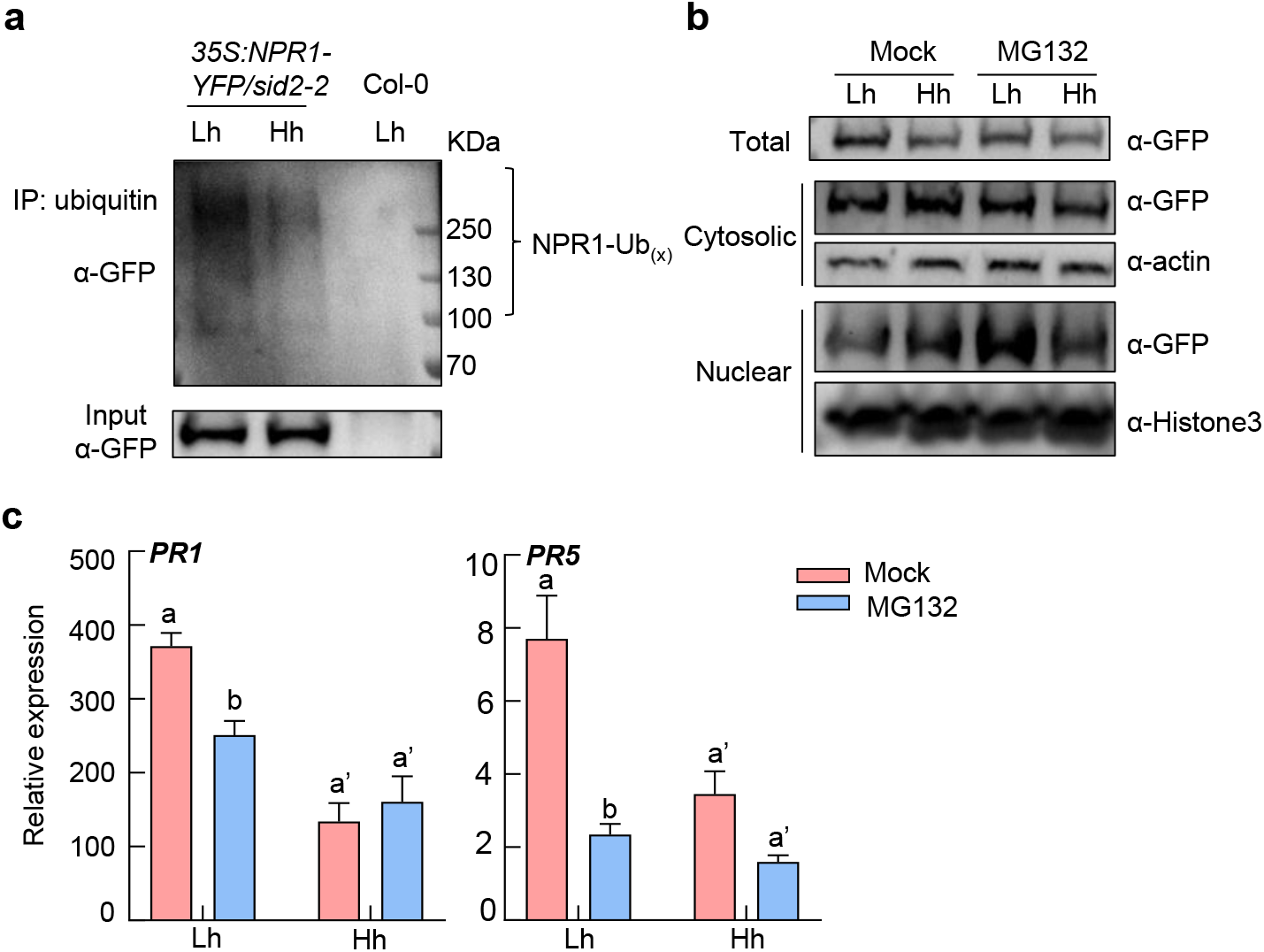
NPR1 poly-ubiquitination is reduced under high humidity and MG132 treatment diminished the high humidity effect on NPR1 protein level and SA gene expression in the *35S:NPR1-YFP/sid2-2* plants. **a**, The *35S:NPR1-YFP/sid2-2* plants were pre-treated with Lh or Hh for 24 h, infiltrated with 50 μM MG132 solution, air-dried for ~0.5 h, and then sprayed with 100 μM BTH. Samples were taken 12 h after BTH treatment. Ubiquitinated proteins were enriched using anti-ubiquitin beads. NPR1-GFP (NPR1-Ubx) in input and IP samples were detected by western blot with anti-GFP antibody. **b-c**, The plants were treated the same as in **a**. **b**, Total, cytosolic and nuclear proteins were extracted and subject to SDS-PAGE and western blot. NPR1-YFP protein was detected by anti-GFP antibody. Actin and Histone3 were used as loading controls. **c**, The *PR* gene expression was analyzed by RT-qPCR. Data represent mean ± SEM (n = 3 biological replicates). Different letters indicate statistically significant differences, as analyzed by two-way ANOVA with Tukey’s test (p<0.05). Experiments were repeated three times with similar results.

**Extended Data Fig. 8.**
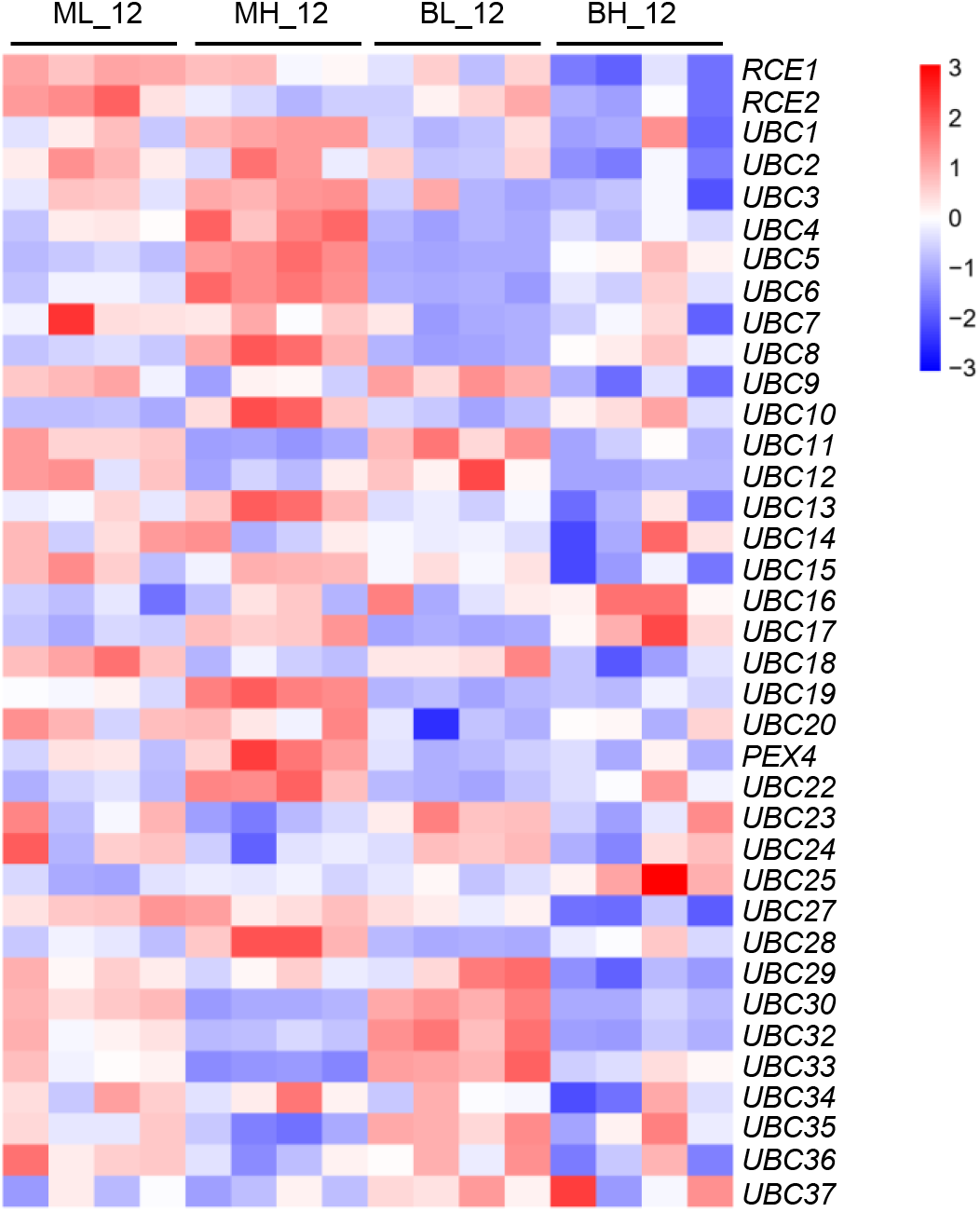
The genes encoding E2 ubiquitin conjugating enzymes in Arabidopsis are not significantly down-regulated under high humidity, based on our RNAseq analysis. A heatmap based on the TPM of each E2 gene is shown.

**Extended Data Fig. 9.**
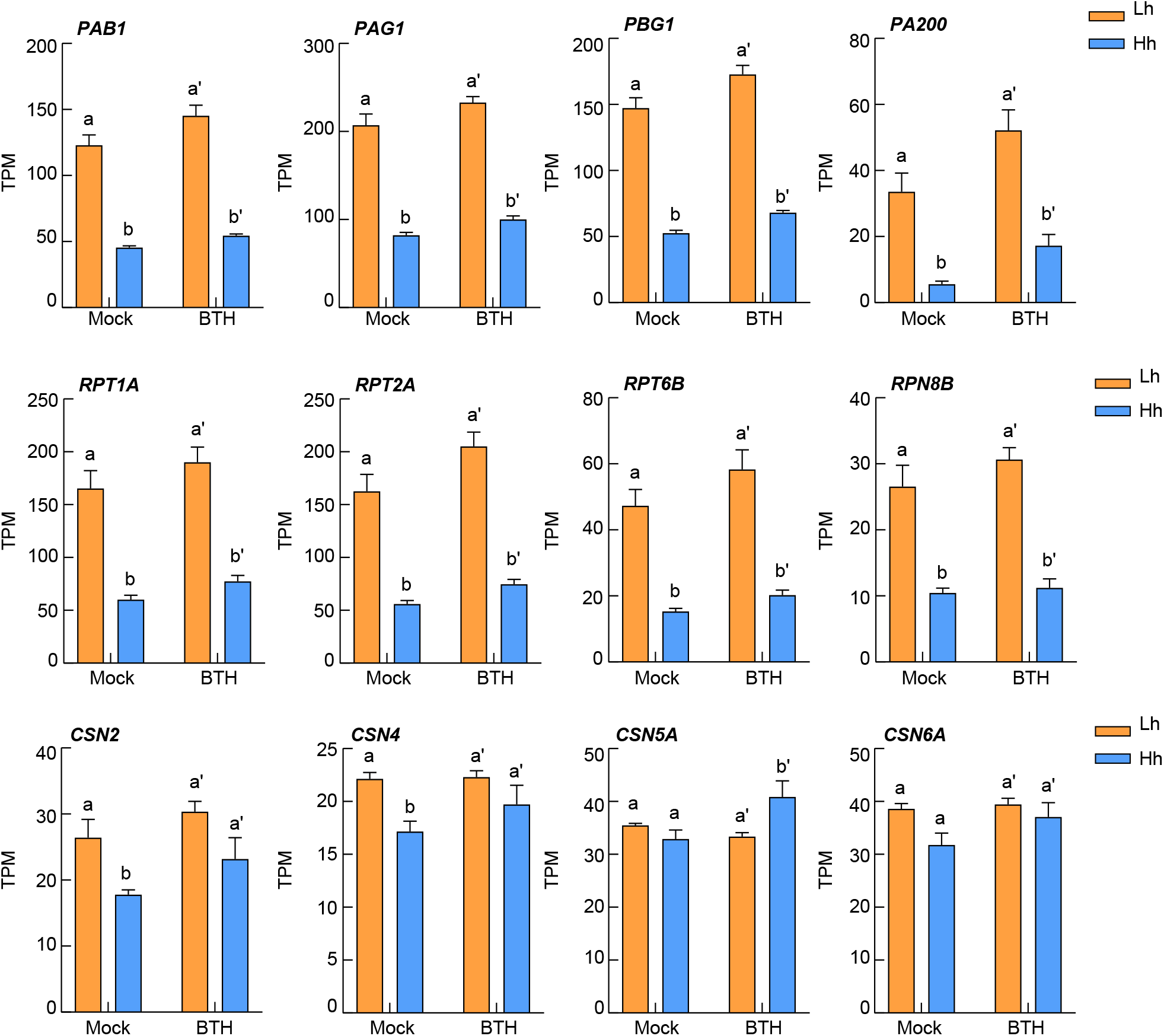
The 26S proteasome machinery genes are down-regulated, but the COP9 signalosome genes are not significantly affected under high humidity, based on our RNAseq analysis. TPMs of representative genes of the 26S proteasome machinery (*PAs/PBs/RPTs/RPNs/PA200*) and COP9 signalosome (CSNs) are shown. Data represent mean ± SEM (n = 4 biological replicates). Different letters indicate statistically significant differences, as analyzed by two-way ANOVA with Tukey’s test (p<0.05).

**Extended Data Fig. 10.**
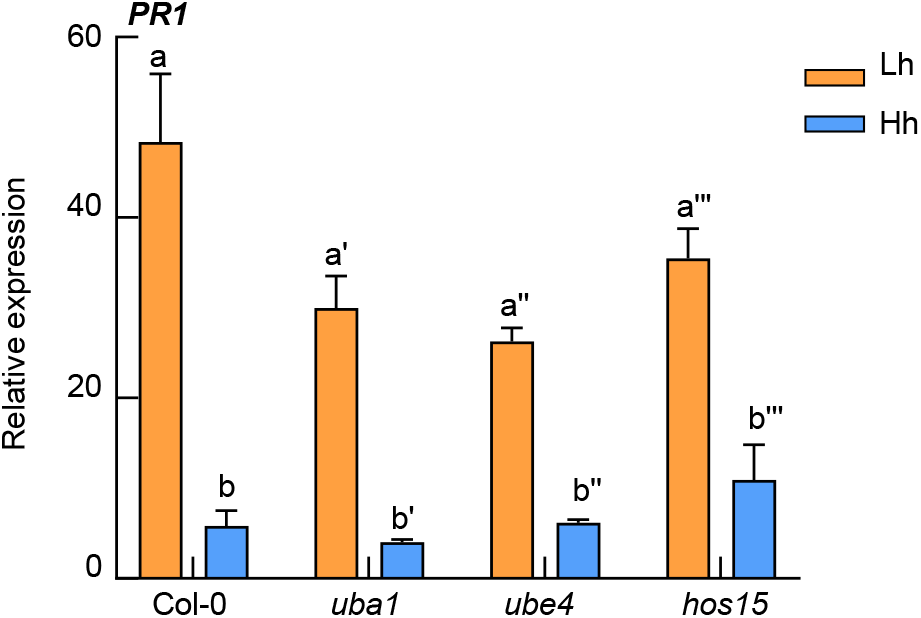
RT-qPCR analysis of *PR1* expression level in Col-0, *uba1* (E1), *hos15* (E3) and *ube4* (E4) mutant plants. Plants were pre-treated with different humidity for 24 h, sprayed with 100 μM BTH and sampled 12 h after BTH treatment. Data represent mean ± SEM (n = 3 biological replicates). Different letters indicate statistically significant differences, as analyzed by two-way ANOVA with Tukey’s test (p<0.05). Experiments were repeated three times with similar results.

## Notes

https://www.ncbi.nlm.nih.gov/geo/

### Competing Interest Statement

The authors have declared no competing interest.

